# EMP3 sustains oncogenic EGFR/CDK2 signaling by restricting receptor degradation in glioblastoma

**DOI:** 10.1101/2023.05.06.539678

**Authors:** Antoni Andreu Martija, Alexandra Krauß, Natalie Bächle, Laura Doth, Arne Christians, Damir Krunic, Martin Schneider, Dominic Helm, Rainer Will, Christian Hartmann, Christel Herold-Mende, Andreas von Deimling, Stefan Pusch

**Affiliations:** Clinical Cooperation Unit (CCU) Neuropathology, German Cancer Research Consortium (DKTK), German Cancer Research Center (DKFZ), Heidelberg, Germany; Department of Neuropathology, Heidelberg University Hospital, Heidelberg, Germany; Faculty of Biosciences, Heidelberg University, Heidelberg, Germany; Faculty of Medicine, Heidelberg University, Heidelberg, Germany; Present address: Institute for Molecular Medicine, University Medical Center of the Johannes Gutenberg-University Mainz, Mainz, Germany; Department of Neuropathology, Institute of Pathology, Hannover Medical School, Hannover, Germany; Present address: Canopy Biosciences, Bruker Nano Group, Hannover, Germany; Light Microscopy Facility, DKFZ, Heidelberg, Germany; Mass Spectrometry-Based Protein Analysis Unit, Genomics and Proteomics Core Facility, DKFZ, Heidelberg, Germany; Cellular Tools/Vector & Clone Repository Unit, Genomics and Proteomics Core Facility, DKFZ, Heidelberg, Germany; Department of Neurosurgery, Heidelberg University Hospital, Heidelberg, Germany

**Keywords:** CDK2, EGFR, EMP3, TBC1D5, glioblastoma

## Abstract

Epithelial membrane protein 3 (EMP3) is an N-glycosylated tetraspanin with a putative trafficking function. It is highly expressed in isocitrate dehydrogenase-wild-type glioblastoma (IDH-wt GBM), and its high expression correlates with poor survival. However, the exact trafficking role of EMP3 and how it promotes oncogenic signaling in GBM remain unclear. Here, we show that EMP3 promotes EGFR/CDK2 signaling by regulating the trafficking and enhancing the stability of EGFR. BioID2-based proximity labeling revealed that EMP3 interacts with endocytic proteins involved in the vesicular transport of EGFR. EMP3 knockout (KO) enhances epidermal growth factor (EGF)-induced shuttling of EGFR into RAB7+ late endosomes, thereby promoting EGFR degradation. Increased EGFR degradation is rescued by the RAB7 negative regulator and novel EMP3 interactor TBC1D5. Phosphoproteomic and transcriptomic analyses further showed that EMP3 KO converges into the inhibition of the cyclin-dependent kinase CDK2 and the repression of EGFR-dependent and cell cycle transcriptional programs. Phenotypically, EMP3 KO cells exhibit reduced proliferation rates, blunted mitogenic response to EGF, and increased sensitivity to the pan-kinase inhibitor staurosporine and the EGFR-specific inhibitor osimertinib. Furthermore, EGFR-dependent patient-derived glioblastoma stem cells display increased susceptibility to CDK2 inhibition upon EMP3 knockdown. Lastly, using TCGA data, we showed that GBM tumors with high EMP3 expression have increased total and phosphorylated EGFR levels. Collectively, our findings demonstrate a novel EMP3-dependent mechanism by which EGFR/CDK2 activity is sustained in GBM. Consequently, this stabilizing effect provides an additional layer of tumor cell resistance against targeted kinase inhibition.

## Introduction

Epithelial membrane protein 3 (EMP3) is an *N-*glycosylated tetraspanin implicated in isocitrate dehydrogenase-wild-type (IDH-wt) glioblastoma (GBM) [10, 11, 27, 39]. EMP3 has low expression in the adult brain but is highly expressed in IDH-wt GBM [27, 39, 62]. High EMP3 expression is positively associated with poor survival of GBM patients [12, 16, 18, 27, 62]. Studies have shown that EMP3 supports TGF-β signaling in CD44-high GBM cells [27] and induces an immunosuppressive GBM microenvironment characterized by reduced T cell infiltration and increased amounts of M2 macrophages in animal models [10]. In non-glioma cells, EMP3 facilitates the activation of receptor tyrosine kinase (RTK) signaling pathway components, including the epidermal growth factor receptor (EGFR) [11, 21, 59]. This is proposed to occur through EMP3’s putative trafficking function, after a yeast two-hybrid screen identified EMP3’s interaction with several EGFR trafficking regulators [11]. However, these mechanisms, along with their potential downstream consequences, have not been further elucidated. To date, whether and how EMP3 regulates EGFR trafficking and signaling in IDH-wt GBM are still unclear.

Aberrant EGFR activation is frequently observed in IDH-wt GBM [37, 40]. This usually results from *EGFR* gene amplification or from mutations that promote constitutive activation of the receptor [22, 43]. Once activated, EGFR undergoes autophosphorylation, thereby stimulating downstream signaling most notably through the PI3K/AKT and MAPK pathways [22]. In non-malignant cells, this signaling cascade is controlled by several homeostatic mechanisms, including endolysosomal degradation of EGFR [1, 8]. After binding to EGF, phosphorylated EGFR is internalized into endosomes and eventually shuttled into lysosomes, leading to receptor degradation and termination of signaling [1]. However, tumor cells can bypass this process by inhibiting receptor internalization, promoting receptor recycling, or attenuating receptor degradation [8, 22, 41].

In this study, we integrated protein-protein interaction screens, phosphoproteomics, transcriptomics, and loss of function studies to uncover how EMP3 regulates EGFR trafficking and signaling in IDH-wt GBM cells. Loss of EMP3 increases EGFR trafficking towards RAB7^+^ late endosomes and EGF-induced degradation of EGFR. Enhanced EGFR degradation is rescued by overexpression of the novel EMP3 interactor TBC1D5, a GTPase-activating protein (GAP) and negative regulator of RAB7. Consistent with compromised EGFR function, EMP3 KO inactivates the EGFR effector CDK2 and inhibits transcription of EGFR-responsive cell cycle genes. Moreover, loss of EMP3 led to decreased proliferation, reduced mitogenic response to EGF, and increased susceptibility to pan-kinase and targeted inhibition of EGFR and CDK2. Lastly, analysis of The Cancer Genome Atlas (TCGA) dataset showed that GBM tumors with high EMP3 expression have higher levels of total and phosphorylated EGFR. Taken together, our findings demonstrate that EMP3 supports EGFR/CDK2 signaling and confers IDH-wt GBM cells with therapeutic resistance against kinase inhibition.

## Materials and Methods

### Cell culture

U-118 and DK-MG cells were purchased from the American Type Culture Collection (ATCC, Virginia, USA) and the German Collection of Microorganisms and Cell Culture (DSMZ, Braunschweig, Germany), respectively. Both cell lines were authenticated by Multiplexion (Heidelberg, Germany) and checked for mycoplasma contamination by GATC (Ebersberg, Germany). Both cell lines were grown as adherent monolayers and maintained at 37°C, 5% CO_2_ in Dulbecco’s Modified Eagle Medium (DMEM) with high glucose, GlutaMAX and pyruvate (Gibco 31966047) supplemented with 10% Fetal Bovine Serum (Gibco 10500-064) and 1% antibiotic-antimycotic (Gibco 15240-062). The glioblastoma stem cell (GSC) lines NCH1425 and NCH644 were obtained from the Department of Neurosurgery in Heidelberg University Hospital. GSCs were grown as tumor spheroids and maintained in DMEM/F-12 with GlutaMAX (Gibco 31331028) supplemented with B-27 (Gibco 12587010), 20 ng/mL human EGF (Peprotech AF-100-15-100), 20 ng/mL of human FGF-basic (Peprotech 100-18B-250) and 1% antibiotic-antimycotic. Stably transfected or lentivirally transduced cells were selected and maintained in the same medium supplemented with 0.5 to 1 µg/mL puromycin (MP Biomedicals 194539) or 4 µg/mL blasticidin (US Biological Life Sciences B2104-30). Generation of CRISPR/Cas9-edited EMP3 KO cells and other cell lines stably expressing cloned plasmids are further described in the Supplementary Materials and Methods.

### BioID2-based proximity labeling

Total cell lysates were collected from U-118 cells expressing BioID2 bait constructs and treated with 50 µM biotin (Sigma-Aldrich B4639) for 18 hours. Lysates were subjected to streptavidin pull-downs using Pierce High Capacity Streptavidin Agarose Resin (Thermo Fisher Scientific 20357) following the manufacturer’s protocol. Eluates were then subjected to liquid chromatography – tandem mass spectrometry (LC-MS/MS) analysis using the Ultimate 3000 UPLC system directly connected to an Orbitrap Exploris 480 mass spectrometer (Thermo Fisher Scientific). Protein identification and quantification were carried out using MaxQuant version 1.6.14.0 [55] and statistical analysis was performed using Perseus 1.6.14.0 [56]. Network and data visualization were further performed using Cytoscape version 3.9.1 [50] and ProHits-viz [29]. Sample preparation as well as data acquisition and analysis settings are further detailed in the Supplementary Materials and Methods.

### Phosphoproteomic analysis

Total cell lysates were collected from GBM control and EMP3 KO cells lysed with 1% sodium dodecyl sulfate and 1% sodium deoxycholate in 100 mM triethylammonium bicarbonate (TEAB; Thermo Fisher Scientific PI90114) containing protease and phosphatase inhibitors. Lysates were heated at 95°C for 5 minutes, sonicated for 5 cycles (35% power for 20 seconds per cycle), and clarified by centrifugation at 20,000 x g at 4°C for 10 minutes. Proteins were precipitated using chloroform/methanol as previously described [60]. Protein pellets were then resuspended (8 M Urea, 100 mM NaCl, 50 mM TEAB, pH 8.5), followed by reduction in 10 mM DTT for 1 hour at 27°C, alkylation by 30 mM iodoacetamide for 30 minutes at room temperature in the dark and quenching the reaction by adding additional 10 mM DTT. Samples were subsequently digested by Lys-C at an enzyme:protein ratio of 1:100 for 3-4 hours at 30°C, diluted with 50 mM TEAB to achieve a final urea concentration of 1.6 M, and further digested with trypsin overnight at 37°C in an enzyme:protein ratio of 1:50. Digestion was stopped by acidification using TFA (0.02% (vol/vol) TFA). Digested peptides were desalted using C18 SepPack cartridges and resuspended in 0.07 % (v/v) TFA in 30 % (v/v) ACN and fractionated by on-column Fe^3+^- IMAC enrichment on an Ultimate 300 LC system using a previously described method [46]. The two resulting fractions per sample, containing either unphosphorylated or phosphorylated peptides, were desalted by StageTips [45]. Prior to LC-MS/MS analysis the dry peptides were resolved in 50 mM citric acid and 0.1 % TFA. LC-MS/MS analysis was performed using the mass spectrometer described above and following the settings described in the Supplementary Materials and Methods. Protein quantification and statistical analysis were performed using MaxQuant version 1.6.14.0 [55]. Phosphorylation analysis was performed using Ingenuity Pathway Analysis version 22.0 (Qiagen). Kinase enrichment analysis and upstream kinase prediction were additionally performed using Kinase Enrichment Analysis version 3 [32] and Robust Inference of Kinase Activity version 2.1.3 [61], respectively. Additional details about data acquisition and analysis are further detailed in the Supplementary Materials and Methods.

### Microarray analysis

Total RNA were extracted from GBM control and EMP3 KO cells using the Nucleospin RNA kit (Macherey-Nagel) following the manufacturer’s instructions. Samples were sent to the Microarray Unit of the DKFZ Genomics and Proteomics Core Facility (Heidelberg, Germany), which performed quality control followed by microarray hybridization using the Human Clariom S assay (Applied Biosystems). To analyze microarray data, raw CEL files were imported into the Transcriptome Analysis Console software version 4.0.2.15 (Thermo Fisher Scientific). Differentially expressed genes (DEGs) between control and EMP3 KO cells were identified by filtering for genes with a linear fold-change ≤ -2 (i.e., downregulated) or ≥ 2 (i.e., upregulated) and false discovery rate (FDR)-adjusted p-value ≤ 0.05. DEG lists were then imported into the Ingenuity Pathway Analysis software version 22.0 (IPA, Qiagen). Core analysis was performed on all downregulated and upregulated genes using the default IPA settings, except for the Species parameter, which was restricted to “Human” only. KEGG pathway analysis of DEGs and master regulators was also performed using the Cytoscape stringAPP plug-in (version 1.7.1). Gene set enrichment analysis was also performed using GSEA version 4.2.3 [53]. KEGG pathway analysis of DEGs and master regulators was also performed using the Cytoscape stringAPP plug-in (version 1.7.1). qPCR validation of selected DEGs was performed as described in the Supplementary Materials and Methods.

### Proximity ligation assay

Proximity ligation assays (PLA) were performed using the NaveniFlexMR PLA kit (Navinci). Briefly, cells seeded in 8-well chamber slides were fixed with 4% formaldehyde for 15 minutes and permeabilized with 0.1% Triton-X in PBS for 10 mins after 3 PBS washes. Afterwards, fixed cells were washed with PBS thrice then incubated with blocking buffer at 37°C for 1 hour. Primary antibody incubation was then performed overnight at 4°C using the appropriate antibody dilutions. The next day, cells were incubated with the anti-mouse and anti-rabbit Navenibodies (Navinci) at 37°C for 1 hour. Afterwards, the three enzymatic reactions were carried out following the manufacturer‘s protocol. After the third reaction, cells were washed in 1X TBS for 10 mins twice, then finally washed in 0.1X TBS for 15 mins. Slides were then dried, mounted with VectaShield® HardSet™ with DAPI, and covered with coverslips. Images were captured as specified above. For each field, PLA signals were measured and normalized to nuclei count using an in-house ImageJ script provided by the Light Microscopy Facility of the DKFZ.

### Measurement of EGFR activation and degradation kinetics

A total of 2.0 x 10^5^ GBM cells in 2 mL of DMEM maintenance medium were seeded into 35- mm cell culture dishes and were incubated at 37°C, 5% CO_2_ overnight. After overnight incubation, cells were serum-starved by replacing old medium with an equivalent volume of DMEM and 1% antibiotic-antimycotic. Serum-starved cells were incubated overnight. Cells were then treated with 100 µg/mL of the protein synthesis inhibitor cycloheximide (Sigma-Aldrich C4859) for 1 hour. Afterwards, the old serum-free medium was replaced with new serum-free media with or without 100 ng/mL EGF (Peprotech AF-100-15). For each experimental condition (i.e., control and EMP3 KO), four dishes containing EGF-treated cells were incubated for 2 hours, and lysates were collected at the following intervals: t = 30, 60, 90, and 120 min after EGF treatment. Untreated cells served as control and was considered as time point t = 0. Lysates were stored in -80°C until Western blot was performed.

### Cell proliferation and apoptosis assays

Cell proliferation was measured using CellTiter-Glo 3D (Promega G9683) following the manufacturer’s instructions. Caspase 3/7 activity was measured using the Caspase-Glo 3/7 Assay Kit (Promega G8091). Pre-treatment of cells with EGF, staurosporine, AZD9291, and K03861, as well as measurement of apoptotic activity via PARP cleavage, are further described in the Supplementary Materials and Methods.

### Statistical analyses

All other data apart from the LC-MS/MS data were statistically analyzed in GraphPad Prism version 9.3.1. Unpaired t-test was used when comparing two groups with equal variances. Welch’s t-test was used when comparing two groups with unequal variances. Welch’s ANOVA with Dunnett’s T3 multiple comparisons test was used when comparing more than two groups with unequal variances.

## Results

### Establishment of BioID2-based proximity labeling in GBM cells

To define EMP3’s interactome, we performed BioID2-based proximity labeling [28] using U- 118 GBM cells (Supplementary Fig. S1A-C). We stably transfected U-118 cells with pMXs plasmids encoding bait proteins tagged with a Myc-glycine-serine linker-BioID2 cassette at their C-terminal ends (Supplementary Fig. S1A). To determine if there are interactions that may be dependent on glycosylation, both wild-type (EMP3 WT) and the glycosylation-deficient N47A mutant forms of EMP3 (EMP3 N47A) were used as baits (Supplementary Fig. S1B). For spatial reference controls, TagRFP and a membrane-localizing form fused to an N-terminal GAP-43-derived membrane-targeting signal (GAP-TagRFP) were used. These controls were selected to identify both cytosolic and plasma membrane interactors of EMP3, since EMP3 resides in both compartments [11, 39, 54]. Immunofluorescence staining (Supplementary Fig. S2A-S2B) and Western blotting of the Myc tag (Supplementary Fig. S2C) confirmed proper subcellular localization and expression of the BioID2-tagged bait proteins. To prepare samples for mass spectrometry analysis, stably transfected cells were treated with 50 µM biotin for 18 hours, and biotinylated proteins were purified by streptavidin pull-down (Supplementary Fig. S1C). Blotting of total lysates with streptavidin-HRP and Coomassie staining verified successful biotinylation and purification of biotinylated proteins, respectively (Supplementary Fig. S2C).

Mass spectrometry analysis of streptavidin pull-downs identified a total of 217 EMP3 WT-proximal proteins when TagRFP was used as the control, and 213 proteins when GAP- TagRFP was used (Fig. 1A). These included proteins that satisfied the filtering criteria (i.e., log2-fold change (FC) vs. spatial reference control ≥ 1; Welch’s t-test p-value ≤ 0.05), as well as proteins that were uniquely detected in the EMP3 WT pull-downs but were absent (i.e., LFQ intensity = 0 in all three replicates) in either spatial reference control (Supplementary File 1). Previously known EMP3 interactors (e.g., CD44, CD47, FLOT1, VAMP3, and ATP5B) were identified using our filtering strategy, thus validating the selected log2-FC and p-value cutoffs (Supplementary Fig. S3A-S3B). Notably, the choice of spatial reference control influenced which proteins were identified as potential EMP3 interactors. Hits that were selectively enriched in EMP3 WT relative to TagRFP cells (EMP3 WT/TagRFP) contained a greater proportion of plasma membrane-localized proteins (52.88% vs. 32.83% with GAP-TagRFP as the control) (Fig. 1B and Supplementary File 2). In contrast, EMP3 WT hits that were enriched relative to GAP-TagRFP (EMP3 WT/GAP-TagRFP) had a bias for cytosolic proteins (42.21% vs. 31.71% in EMP3 WT/TagRFP). Given EMP3’s nature as a trafficking protein, the distinct interactors identified using either the TagRFP or GAP-TagRFP spatial reference control likely represent proteins that preferentially interact with membrane or cytosolic pools of EMP3, respectively. Thus, we took the total pool of EMP3 WT hits consisting of 336 proteins to comprise the full EMP3 interaction network.

**Figure 1.**
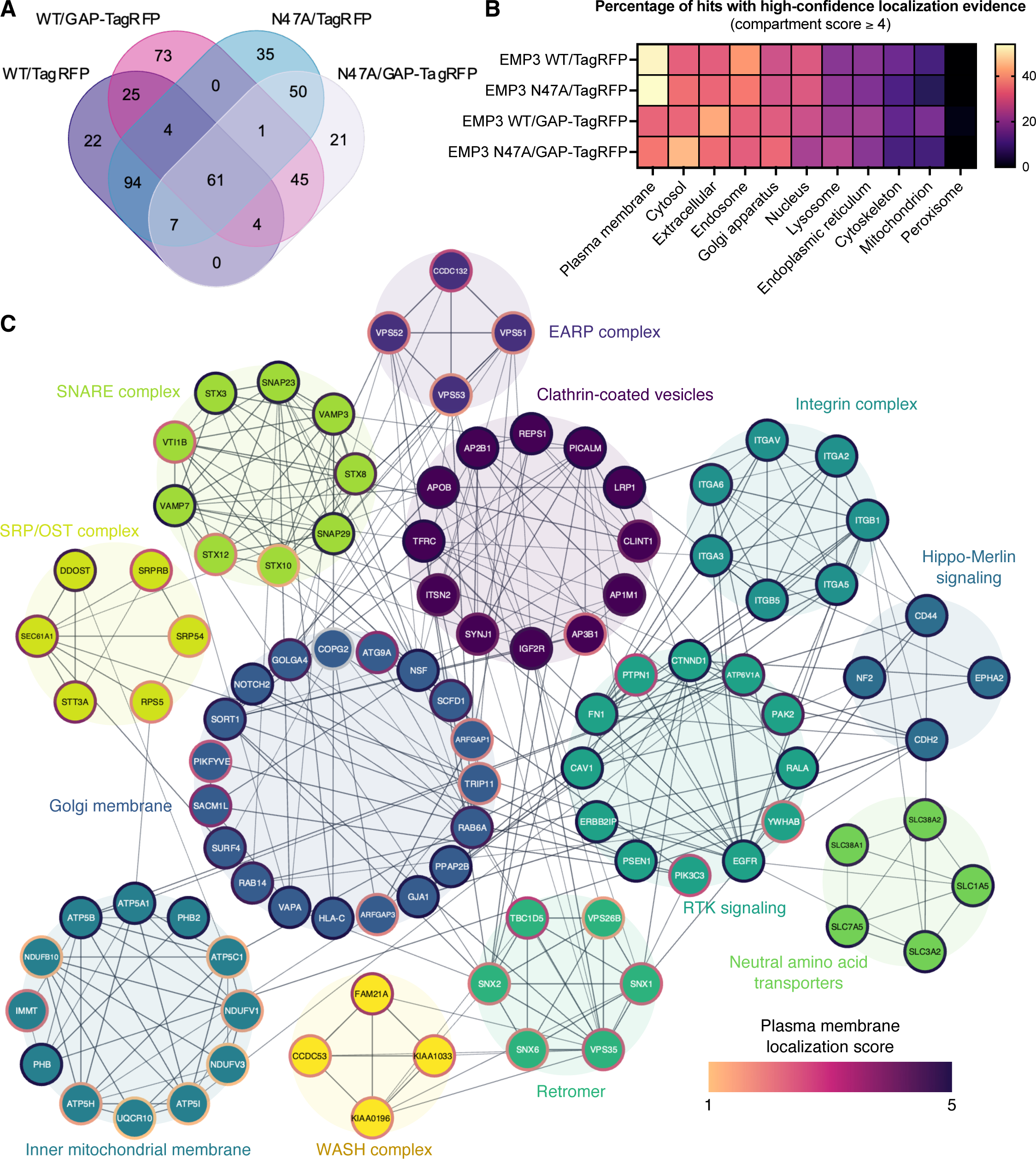
BioID2-based proximity labeling of EMP3 WT and N47A in U-118 glioblastoma cells. **A** Venn diagram showing the number and extent of overlap of identified EMP3-proximal proteins between all pairwise comparisons. **B** Heat map showing the percentage of EMP3-proximal proteins with localization scores ≥ 4 in each cellular compartment and for each pairwise comparison. **C** STRING interaction network resulting from the union of hits identified with EMP3 WT as bait and TagRFP or GAP-TagRFP as spatial reference controls. Edges corresponding to STRING scores > 0.700 and nodes with degrees ≥ 3 were included in the network. Nodes were clustered into functional groups based on STRING enrichment analysis. For simplicity, nodes not belonging to any cluster were removed from the network. Node borders are colored according to plasma membrane localization score, while edge thickness corresponds to STRING interaction scores.

### BioID2-based proximity labeling identifies EGFR and endocytic trafficking regulators as EMP3 interactors

To provide a comprehensive spatial and functional picture of this interactome, network mapping was performed in STRING (version 11.5) using the union of EMP3/TagRFP and EMP3/GAP-TagRFP hits as input. Additional filtering for high-confidence interactions (STRING score ≥ 0.700) and well-connected nodes (degree ≥ 3) followed by enrichment analysis in Cytoscape (version 3.9.1) revealed an interaction network consisting of the GBM driver EGFR, as well as various spatial regulators of receptor trafficking (Fig. 1C). The latter included components of clathrin-coated vesicles (CCVs) as well as retromer, endosome-associated recycling protein (EARP), soluble N-ethylmaleimide-sensitive fusion protein attachment protein receptors (SNARE), and Wiskott Aldrich Syndrome protein and scar homologue (WASH) complexes. Notably, all members of the retromer complex, several CCV-associated proteins (e.g., IGF2R, CLINT1, ITSN2, TFRC, and LRP1), SNARE proteins (e.g., VAMP3; VAMP2, VAMP7, STX12, VTI1B), and the extracellular matrix receptor CD44 were enriched regardless of which spatial reference control was used (Supplementary Fig. S3C). This suggests that these proteins may interact with both membrane or cytosolic pools of EMP3 or consistently co-traffic with EMP3 as it moves to and from both cellular compartments. Expectedly, most transmembrane proteins were preferentially enriched in the EMP3/TagRFP set and therefore were more likely to be proximal to membrane pools of EMP3. The EARP complex, which is involved in recycling internalized receptors back to the cell surface [47], was also selectively enriched in this set. Proximity ligation assay (PLA) in U-118 cells further validated EGFR, CLINT1, the EARP component VPS53, and several retromer proteins (TBC1D5, SNX1, SNX2) as EMP3 interactors (Fig. 2).

**Figure 2.**
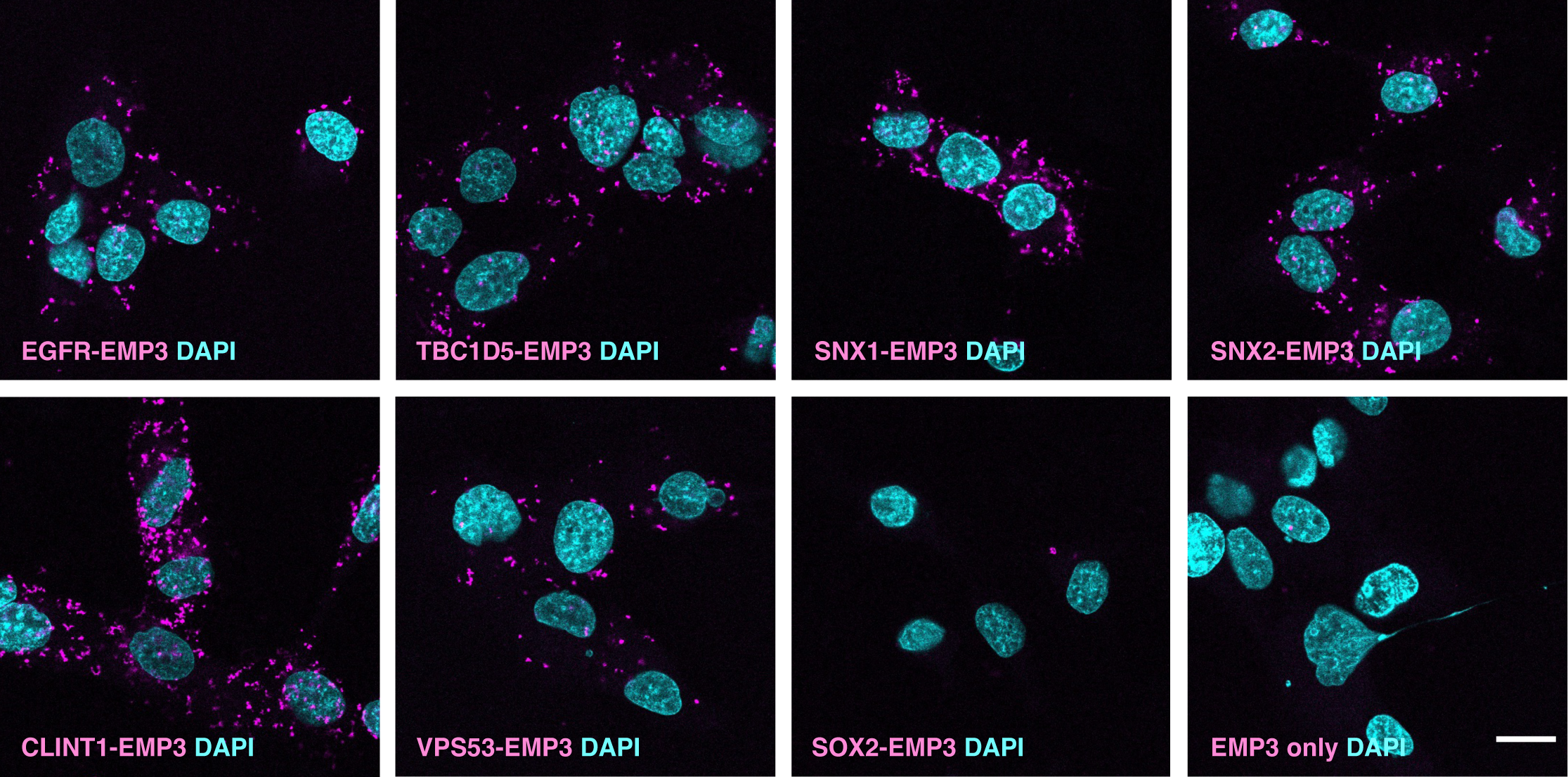
Validation of selected BioID2-identified EMP3-proximal proteins using proximity ligation assay (PLA). PLA signals (magenta) indicating physical proximity (∼40 nm) between EMP3 and various BioID2 hits, including EGFR, retromer components (TBC1D5, SNX1, SNX2), clathrin- coated vesicle-associated protein CLINT1 and EARP complex member VPS53. SOX2 was used as negative control. Nuclei are colored in cyan. Scale bar = 20 μm.

On the other hand, the cytosolic pool of EMP3 selectively interacted with proteins that facilitate targeting of nascent multi-pass membrane proteins towards the ER membrane (e.g., SRPRB, SRP54, SEC61A1). Cytosolic EMP3 was also selectively proximal to components of the oligosaccharyltransferase (OST) complex (e.g., STT3A, DDOST), which facilitate co-translational *N-*glycosylation of nascent proteins [20, 51]. Interestingly, the interaction with DDOST was lost when the EMP3 N47A mutant was used as bait, hinting that the asparagine residue at this position may mediate proper recognition of this OST subunit. We also observed several Golgi- and inner mitochondrial membrane-localizing hits that were not significantly enriched when EMP3 N47A was used as bait (Supplementary Fig. S3C), suggesting that glycosylated EMP3 may uniquely localize toward these compartments. Nonetheless, most membrane and trafficking proteins retained their enrichment even with the N47A mutation (Supplementary Fig. S3C). Collectively, these results indicate that while *N*-linked glycosylation may confer EMP3 with an organelle-specific function, it is not required for its membrane localization and interaction with most trafficking regulators.

### EMP3 restricts EGF-induced late endosomal shuttling of EGFR and its eventual degradation in a TBC1D5-dependent manner

Given its association with membrane receptors and trafficking regulators, we then hypothesized that EMP3 may actively regulate certain receptor trafficking events. Thus, we focused our investigations on the possible effect of EMP3 on the intracellular trafficking of EGFR. To this end, we first generated and validated CRISPR/Cas9-edited EMP3 knockouts using U-118 and DK-MG GBM cells (Supplementary Fig. S4 A-C). Afterwards, we examined the kinetics of EGFR activation and degradation upon EGF treatment of control and EMP3 KO cells pre-treated with 100 µg/mL of the protein synthesis inhibitor cycloheximide for 1 hour. Apart from inducing EGFR phosphorylation, EGF also stimulates the internalization and subsequent degradation of EGFR in lysosomes [1]. Consistent with this, Western blots showed continuous EGFR degradation in EGF-treated cells over the course of 2 hours. Noticeably, ligand-induced degradation of total and phosphorylated EGFR (Tyr 1068) was accelerated in EMP3 KO cells (Fig. 3A and 3B), indicating that EMP3 limits EGF-induced endolysosomal degradation of EGFR. Given EMP3’s interaction with several trafficking proteins, we then hypothesized that EMP3 may attenuate EGFR degradation specifically by restricting receptor trafficking towards degradative endosomes. Supporting this, we observed increased association of phosphorylated EGFR (Tyr1068) with the late endosomal marker RAB7 in EMP3 KO cells 30 minutes after EGF treatment by PLA (Fig. 3C and 3D).

**Figure 3.**
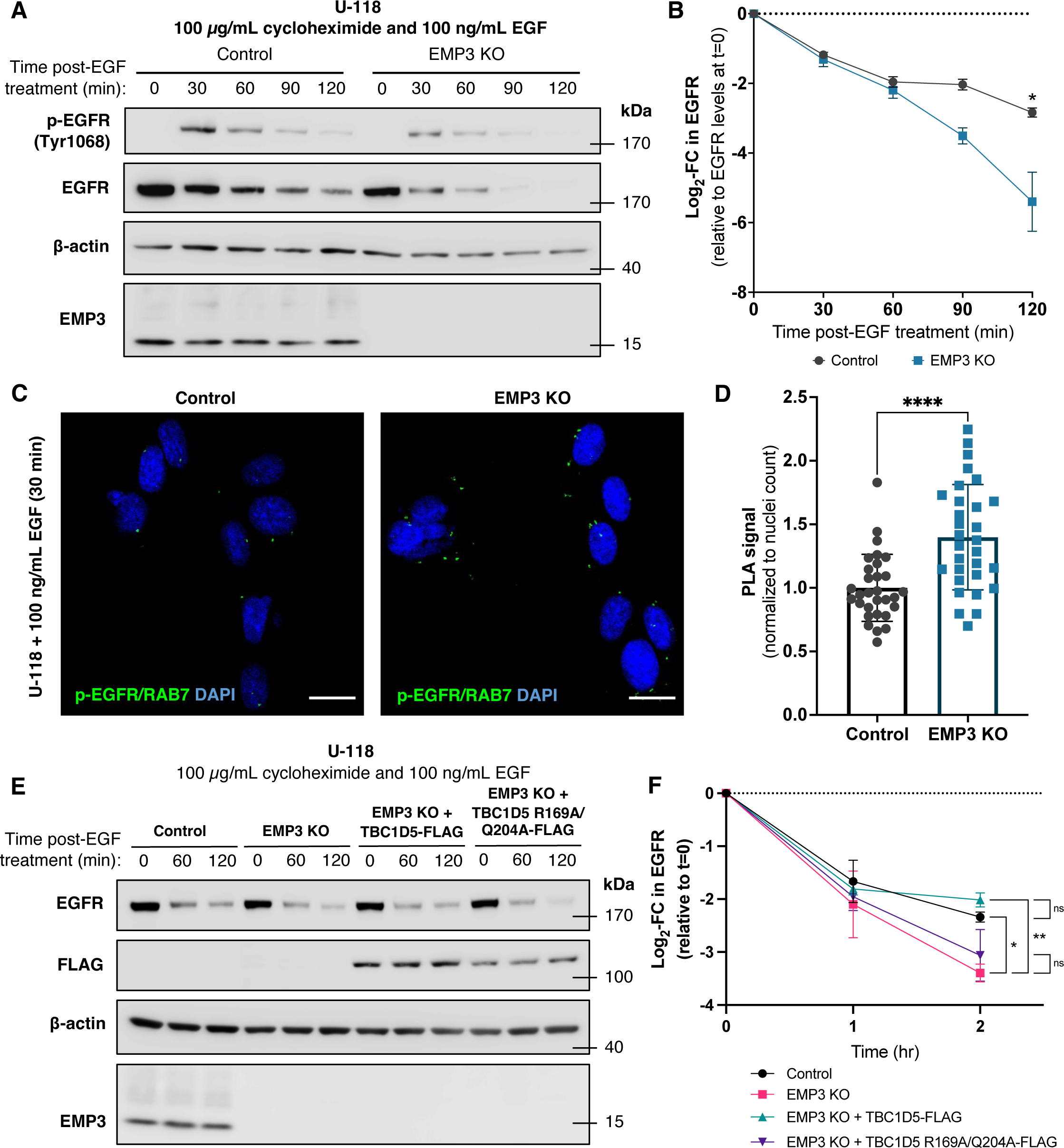
EMP3 restricts RAB7 shuttling and ligand-induced degradation of EGFR in a TBC1D5- dependent manner. **A** Western blot showing the kinetics of EGFR phosphorylation (p-EGFR Tyr1068) and degradation (total EGFR) in U-118 control and EMP3 KO cells over the course of 2 hours after treatment with 100 ng/mL EGF (n=3). **B** Quantification of EGF-induced degradation of EGFR in U-118 control and EMP3 KO cells after treatment with 100 ng/mL EGF (n=3). Band intensities of EGFR were normalized to β-actin, and log2-fold changes (mean + S.E.M.) were calculated relative to the EGFR level at t=0 and plotted over time. (Welch’s two-tailed t-test; * P < 0.05). **C** Representative PLA images testing the association between p- EGFR and RAB7 after 30-min EGF treatment of U-118 control and EMP3 KO cells (n=3). **D** Quantification of p-EGFR-RAB7 PLA signals from EGF-treated U-118 control and EMP3 KO cells. PLA signals were derived from 30 random fields from n=3 independent experiments (10 fields/experiment). Signals were normalized to nuclei count per field (Welch’s two-tailed t-test; **** P = < 0.0001). **E** Western blot showing TBC1D5 WT-mediated rescue of enhanced EGFR degradation in U-118 EMP3 KO cells (n=3). **F** Quantification of EGF-induced EGFR degradation in U-118 control, EMP3 KO, EMP3 KO + TBC1D5 WT, and EMP3 KO + TBC1D5 R169A/Q204A (n=3). Log2-fold changes (mean + S.E.M.) were calculated and plotted as described (Welch’s ANOVA with Dunnett’s T3 multiple comparisons test; * P < 0.05).

To elucidate the mechanism by which EMP3-dependent restriction of EGFR degradation could occur, we focused our attention on TBC1D5, a top interactor of cytosolic EMP3 (Supplementary Fig. S3B) and a retromer component that facilitates GTP hydrolysis and subsequent inactivation of RAB7 [26, 48, 49]. Loss of TBC1D5 function has been shown to convert RAB7 into a hyperactive, GTP-locked state [26]. Active RAB7, in turn, facilitates the fusion of late endosomes with lysosomes, leading to the degradation of late endosomal cargoes like EGFR [1, 3, 14, 23]. Given this information, we hypothesized that EMP3 and TBC1D5 may cooperate to restrict RAB7-mediated degradation of internalized EGFR. Indeed, overexpression of wild-type TBC1D5 (TBC1D5 WT), but not the GAP activity-deficient TBC1D5 R169A/Q204A mutant [19], reversed accelerated degradation of EGFR in EMP3 KO cells (Fig. 3E and 3F). Thus, EMP3 restricts EGFR degradation in a manner that is dependent on the RAB7 negative regulator TBC1D5.

### Phosphoproteomic analysis reveals that EMP3 KO converges into CDK2 inhibition

To identify what signaling defects result from the loss of EMP3 and its EGFR-stabilizing effect, we performed mass spectrometry-based phosphoproteomic analysis of EMP3 KO and control cells cultured in normal maintenance medium for 72 hours. On average, we detected a total of 8892 class I serine/threonine/tyrosine (STY) phosphosites (i.e., localization probability ≥ 0.75) across all samples, while an average of 4013 proteins were quantified in a full proteome analysis performed in parallel (Supplementary Fig. S5A-S5B). For downstream analysis, we only considered phosphosites that were detected in at least 2 out of 3 replicates per condition. Using this filter, we identified 1408 differentially phosphorylated proteins (i.e., proteins with phosphosite |log2-FC| between EMP3 KO vs. control ≥ 1; FDR-adjusted p-value ≤ 0.05) in DK- MG cells (Supplementary File 3). In U-118 cells, there were 435 differentially phosphorylated proteins between EMP3 KOs and controls (Supplementary File 3). Majority of the differentially phosphorylated proteins were not differentially abundant at the full proteome level, indicating that most of the phosphorylation changes were driven by signaling alterations instead of changes in protein abundance. Moreover, both DK-MG and U-118 EMP3 KOs displayed higher percentages of class I phosphosite alterations than protein abundance changes (Supplementary Fig. S5C-S5D). This suggests that globally, EMP3 KO has a greater effect on protein activity than on protein levels. Notably, a higher proportion of DK-MG phosphosites exhibited significant log2-FCs compared to U-118 phosphosites (33.02% vs. 6.98%), indicating that DK-MG is more susceptible to signaling alterations secondary to EMP3 KO (Supplementary Fig. S5C). Intersection of the phosphosites revealed a total of 197 STY residues undergoing phosphorylation changes (i.e., |log2-FC| ≥ 1; FDR-adjusted p-value ≤ 0.05) in both DK-MG and U-118 EMP3 KOs (Fig. 4A). Strikingly, 82.23% of these 197 phosphosites were commonly dephosphorylated (i.e., log2-FC ≤ -1 in both cell lines) upon EMP3 KO. In contrast, only 3.04% phosphosites were commonly phosphorylated (i.e., log2- FC ≥ 1 in both cell lines), while 14.72% were differentially phosphorylated (i.e., log2-FCs in the two cell lines going in opposite directions). Enrichment analysis revealed that the proteins with commonly dephosphorylated sites in the two EMP3 KO cell lines are mostly involved in the cell cycle (Fig. 4B). Thus, EMP3 KO results in the dephosphorylation of cell cycle regulators.

**Figure 4.**
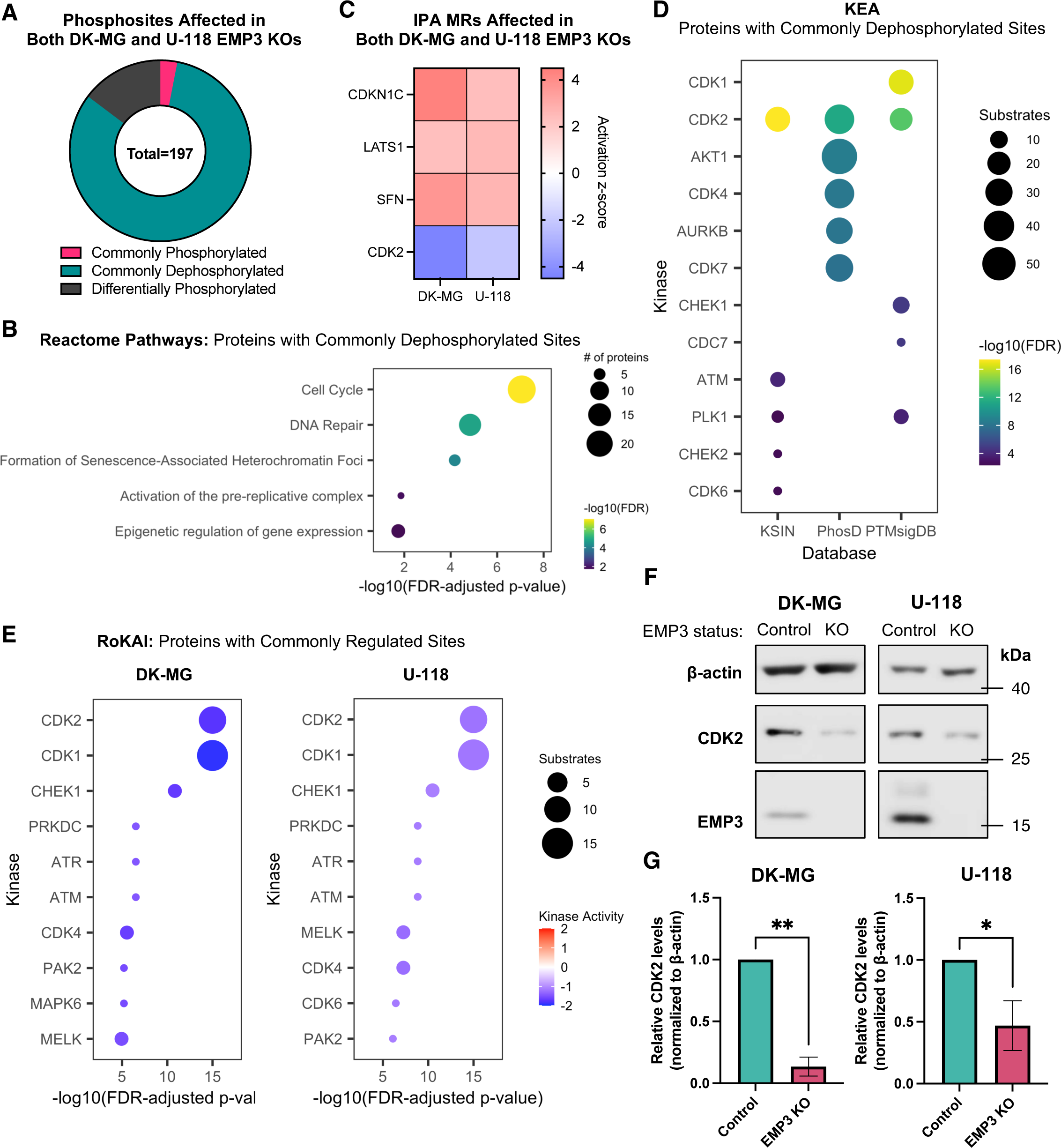
Phosphoproteomic analysis of proteins with commonly regulated phosphosites in DK-MG and U-118 EMP3 KOs. **A** Donut chart showing the distribution of phosphorylation changes common to both DK-MG and U-118 EMP3 KO cells relative to their respective controls. Commonly phosphorylated – log_2_-FC ≥ 1, FDR p-value ≤ 0.05 in both DK-MG and U-118; Commonly dephosphorylated – log_2_-FC ≤ -1, FDR p-value ≤ 0.05 in both DK-MG and U-118; Differentially phosphorylated – FDR p-value ≤ 0.05 but log_2_-FC in opposite directions in DK-MG and U-118. **B** Reactome pathways enriched based on proteins with phosphosites that are dephosphorylated in both DK-MG and U-118 EMP3 KOs. Circle sizes correspond to the number of proteins associated with each term, while the color scale indicates the significance level. **C** Heatmap showing IPA MRs that are activated or inhibited (|activation z-score| ≥ 2, p- value ≤ 0.05) in both DK-MG and U-118 EMP3 KOs and their respective activation z-scores, colored according to predicted activity (red – active; blue – inactive). **D** KEA of proteins with commonly dephosphorylated sites in both DK-MG and U-118 EMP3 KOs. From each kinase- substrate database, the top 5 kinases that phosphorylate substrates in the input list were identified and listed on the y-axis. Circle sizes depict substrate number, while the color scale indicates the significance level of each kinase. **E** RoKAI of proteins with commonly regulated sites in DK-MG (left) and U-118 EMP3 KOs (right). Upstream kinases are listed on the y-axis and ordered according to significance level. Those with -log10(FDR-adjusted p-values) = ∞ were arbitrarily scored as 15 on the x-axis to allow plotting. Circle sizes indicate substrate number, while the color scale shows predicted kinase activity (red – active; blue – inactive). **F** Western blots of CDK2 in control and EMP3 KO cells. **G** Quantification of CDK2 band intensities (mean + S.D.) normalized to β-actin and calibrated relative to control cells (n=3; Welch’s two-tailed t-test; * P < 0.05; ** P < 0.01).

To identify upstream regulators that can explain these phosphoproteomic alterations, we performed phosphorylation analysis using IPA. Intersection of the MRs revealed 4 common alterations in DK-MG and U-118 EMP3 KOs, including inhibition of the cyclin-dependent kinase CDK2 [31] and activation of CDKN1C, a negative regulator of cell proliferation [2] (Fig. 4C). In parallel, we also performed Kinase Enrichment Analysis (KEA) to predict upstream kinases that may be responsible for the observed phosphosite changes. KEA uses multiple kinase-substrate (KS) databases (i.e., KSIN, PhosD, PTMsigDB) to infer upstream kinase regulators from an input list of proteins [32]. Because KEA does not consider the extent of phosphorylation changes (i.e., log2-FC values), we restricted our analysis to proteins with commonly dephosphorylated sites in DK-MG and U-118 EMP3 KOs, as dephosphorylated sites were overrepresented in our analysis. We reasoned that inhibition of certain upstream kinases may account for the high prevalence of these dephosphorylated proteins. KEA consistently predicted CDK2 to be an upstream kinase regulator affected by EMP3 KO across all three KS databases (Fig. 4D). Thus, inhibition of CDK2 activity may largely explain the dephosphorylation of the input substrates in EMP3 KO cells.

One potential limitation of IPA and KEA is that the amino acid identities of the differentially phosphorylated residues are not accounted for in the pathway analysis. To identify upstream kinases based on the phosphosite alterations induced by EMP3 KO, we performed Robust Inference of Kinase Activity or RoKAI [61]. For this analysis, we used commonly regulated STY sites (i.e., FDR-adjusted p-value ≤ 0.05, log2-FC ≥ 1 or ≤ -1 and going in the same direction in both EMP3 KO cell lines) and their corresponding phosphorylation log2-FCs upon EMP3 KO as input. In agreement with the IPA and KEA analyses, RoKAI also indicated significant inhibition of CDK2 based on the phosphorylation changes that occurred in both cell lines (Fig. 4E). To validate our bioinformatic analyses, we assessed and compared the protein levels of CDK2 in control and EMP3 KO cells by Western blotting. Results confirmed that total CDK2 levels are significantly reduced in EMP3 KO cells (Fig. 4F-G). In summary, integration of the three phosphoproteomic analysis pipelines reveals that EMP3 KO converges into CDK2 inhibition and the dephosphorylation of CDK2 substrates involved in cell cycle progression.

### Loss of EMP3 inhibits EGFR-dependent and cell cycle-related transcriptional programs

Next, we sought to determine how loss of EMP3 impacts transcriptional programs. To this end, we performed microarray analysis on total RNA extracted from 72-hour cultures of U-118 and DK-MG cells with or without EMP3. A total of 1191 and 1491 differentially expressed genes (DEGs) were identified upon EMP3 depletion in U-118 and DK-MG cells, respectively (Supplementary File 4). We then performed pathway analyses to identify what transcriptional programs are enriched based on our DEG lists (Fig. 5A-D). KEGG pathway analysis did not reveal any enriched pathways when using the set of 160 genes that were upregulated in both EMP3 KO cell lines. However, consistent with CDK2 inhibition, we identified 125 DNA replication- and cell cycle-related genes that are downregulated in both DK-MG and U-118 EMP3 KOs (Fig. 5A and 5C).

**Figure 5.**
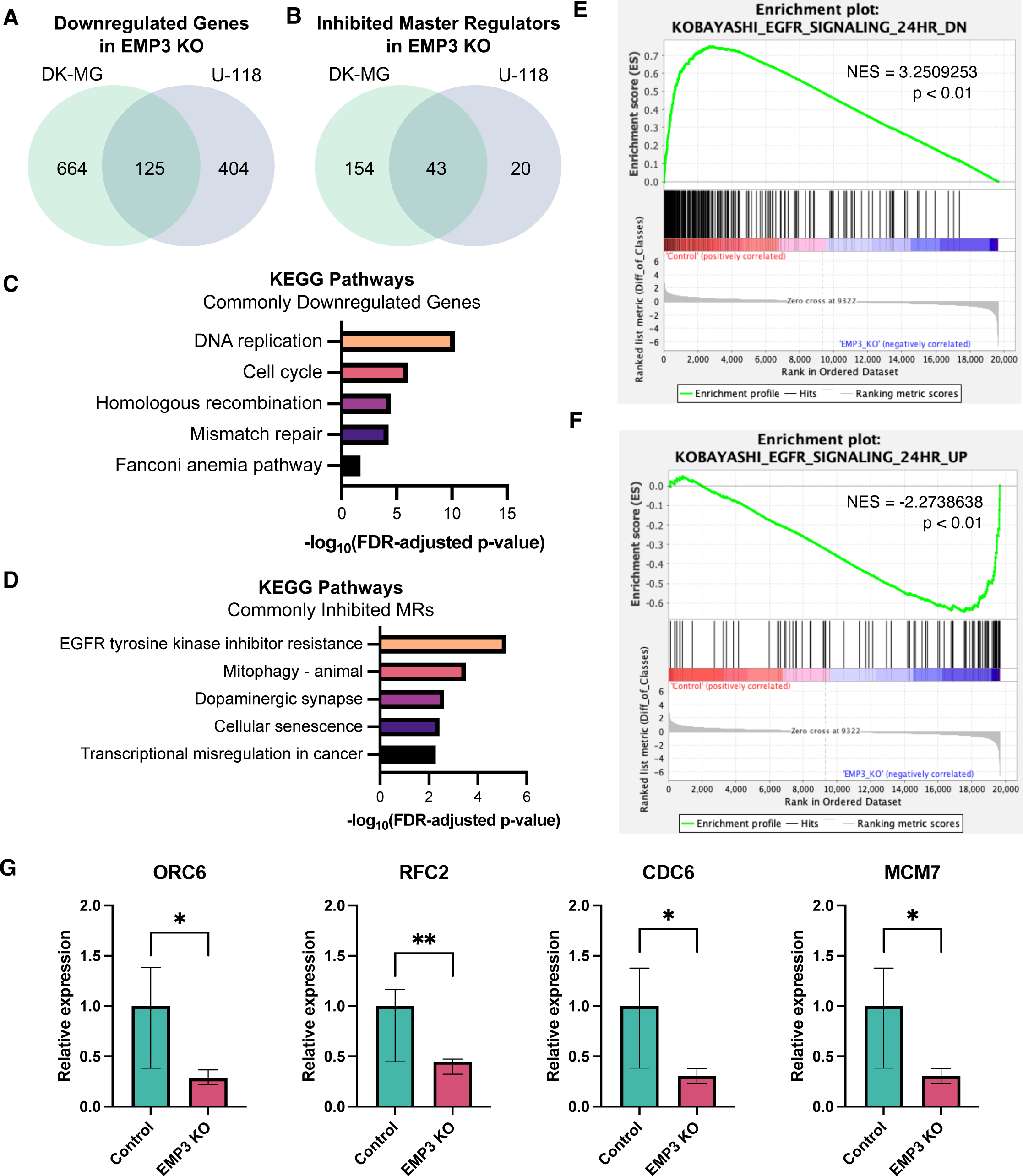
Pathway analysis of differentially expressed genes between control and EMP3 KO cells. **A, B** Venn diagram showing the overlap of (A) downregulated genes (log2-FC ≤ -1, FDR p-value ≤ 0.05) and (B) inhibited master regulators (activation z-score ≤ -2, p-value ≤ 0.05) between DK- MG and U-118 EMP3 KOs. **C, D** KEGG pathways enriched based on the list of (C) commonly downregulated genes and (D) commonly inhibited MRs. **E, F** GSEA results showing upregulation of the gene sets KOBAYASHI_EGFR_SIGNALING_24HR_DN (E) and _UP (F) in U-118 control and EMP3 KO cells, respectively. Genes were sorted from left to right based on the difference of the log2 expression levels between control and EMP3 KO cells. Vertical black bars indicate the location of the genes contributing to the enrichment scores (ES). The ES, which indicate upregulation (ES > 0) or downregulation (ES < 0) of a certain gene, are plotted on the y-axis. NES: normalized enrichment score. **G** qPCR results validating the downregulation of selected genes in U-118 EMP3 KOs (unpaired one-tailed t-test; * P = < 0.05; ** P = < 0.01).

We also used the DEGs from each cell line as input for Ingenuity Pathway Analysis (Qiagen) to predict which upstream signaling pathways are differentially regulated between control and EMP3 KO conditions. This analysis yielded a total of 43 master regulators (MRs) that were inhibited in both DK-MG and U-118 EMP3 KO cells (Fig. 5B). Consistent with impaired EGFR function, EGF was identified to be among the most significantly inhibited MRs in both U-118 (activation z-score = -2.635, P = 2.43 x 10^-32^) and DK-MG (activation z-score = -2.92, P = 1.10 x 10^-17^) EMP3 KO cells (Supplementary Fig. S6A-S6B). KEGG pathway analysis of the 43 commonly inhibited MRs further indicated enrichment of proteins involved in EGFR tyrosine kinase inhibitor resistance (Fig. 5D), hinting that the therapeutic response of GBM cells against EGFR-targeting compounds may be attenuated in the absence of EMP3. In line with this, gene set enrichment analyses (GSEA) of the U-118 and DK-MG DEGs also revealed upregulation of the “KOBAYASHI_EGFR SIGNALING_24HR_DN” gene set in EMP3-expressing control cells (Fig. 5E and Supplementary Fig. S6C). This gene signature includes genes that are downregulated upon EGFR inhibition of non-small cell lung cancer (NSCLC) cells [30]; thus, EMP3 control cells can be presumed to have an intact EGFR function, as these cells retain the expression of genes that are negatively affected by EGFR inhibition. Conversely, U-118 and DK-MG EMP3 KO cells exhibited increased expression of genes that are upregulated upon EGFR inhibition of NSCLC cells (Fig. 5F and Supplementary Fig. S6D). To further validate these transcriptomic results, we selected 4 of the top 5 DEGs (ORC6, RFC2, CDC6, MCM7) that are EGF/EGFR targets according to our IPA and GSEA analyses and are involved in DNA replication or the cell cycle according to KEGG. qPCR analysis confirmed that these 4 genes are downregulated in U-118 and DK-MG EMP3 KOs relative to controls (Fig. 5G and Supplementary Fig. S6E). Taken together, multiple gene enrichment and pathway analyses indicate that EMP3 KO represses the transcription of EGFR-dependent cell cycle genes.

### EMP3 KO reduces mitogenic response to EGF and sensitizes GBM cells to EGFR inhibition

Our functional, transcriptomic, and phosphoproteomic data collectively indicate that targeting EMP3 inhibits EGFR/CDK2 signaling by reducing EGFR stability. To evaluate the phenotypic consequences of this process, we assessed how EMP3 KO impacts the proliferative capacity and response of GBM cells to ligand-dependent EGFR activation. Proliferation rates of EMP3 KO and control cells grown in serum-containing medium were measured for 96 hours. Results consistently showed that in both cell lines, EMP3 KOs were less proliferative than controls (Fig. 6A and 6B). This is consistent with our phosphoproteomics and gene expression data indicating dysregulation of DNA replication and cell cycle progression in EMP3 KOs (Fig. 4B and 5C). To examine the effect of EMP3 on EGFR-dependent proliferation, we serum-starved EMP3 KO and control cells overnight and monitored their proliferation after daily treatment with 100 ng/mL EGF for 72 hours. Serum starvation is presumed to eliminate external sources of mitogenic stimulation; thus, any changes in cell number over this period can be mainly attributed to the externally administered ligand. Consistent with attenuated EGFR signaling, U-118 and DK-MG EMP3 KO cells were less responsive to mitogenic stimulation by EGF (Fig. 6C and 6D).

**Figure 6.**
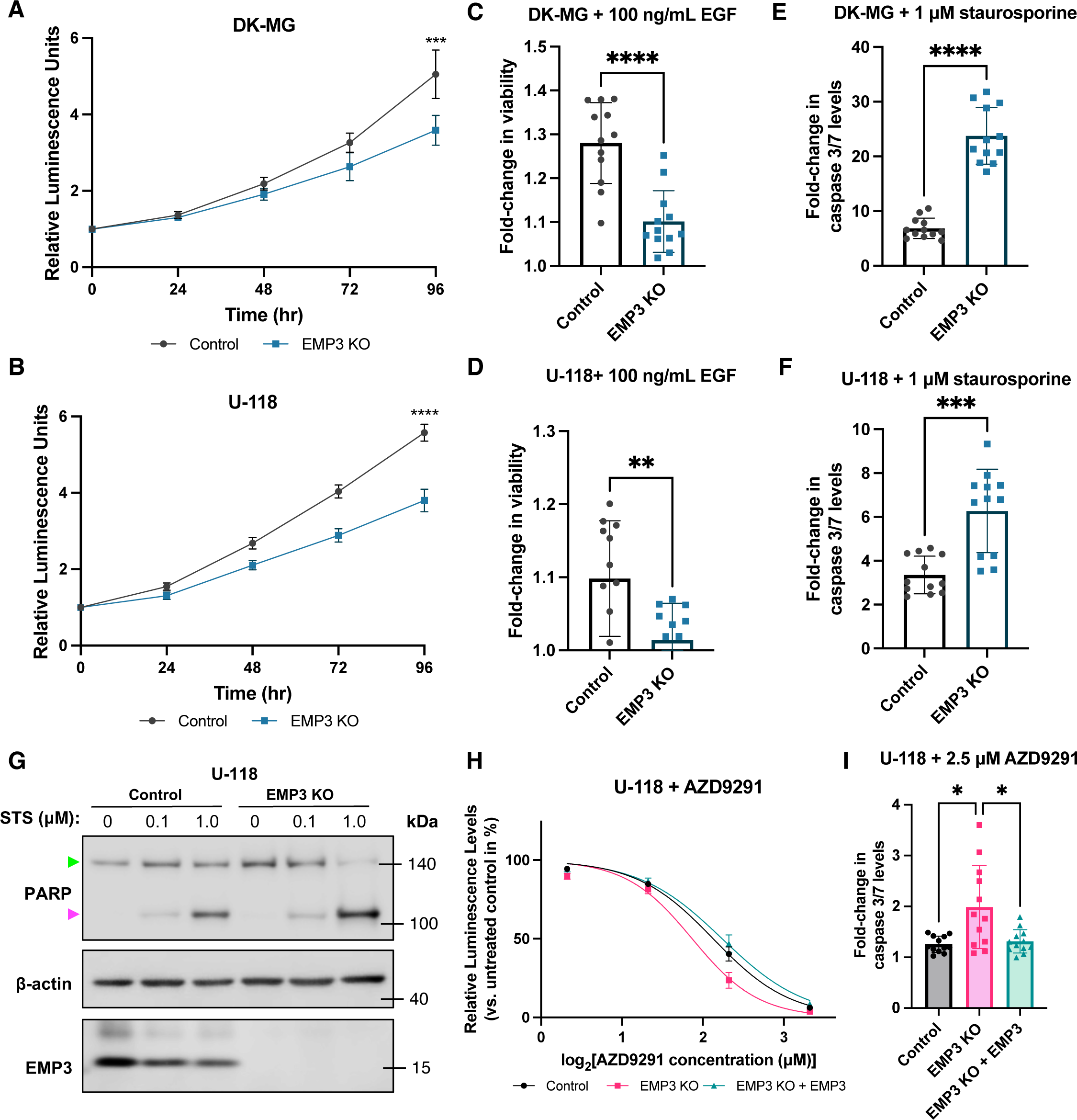
EMP3 KOs have impaired proliferative response and increased sensitivity to kinase inhibition. **A, B** Proliferation of DK-MG (A) and U-118 (B) control and EMP3 KO cells cultured in normal serum-containing medium over the course of 4 days. Dots represent mean fold-changes in CellTiter-Glo luminescence readings relative to t=0, while error bars show 95% confidence intervals (unpaired t-test; ***P < 0.001, ****P < 0.0001). **C, D** Proliferative response of serum- starved DK-MG (C) and U-118 (D) control and EMP3 KO cells to daily EGF treatment for 72 hours. Bar plots represented mean fold-changes in CellTiter-Glo luminescence readings of EGF-treated cells versus untreated controls. Error bars and dots represent S.D. and individual measurements, respectively (unpaired t-test; **P < 0.01, ****P < 0.0001). **E, F** Caspase 3/7 levels in DK-MG (E) and U-118 (F) control and EMP3 KO cells treated with 1 μM staurosporine (STS) for 4 hours. Bar plots represent mean fold-changes in Caspase 3/7 luminescence readings of STS-treated cells versus untreated controls. Error bars and dots represent S.D. and individual measurements, respectively (Welch’s t-test; ***P < 0.001, ****P < 0.0001). **G** Western blot showing full-length (green arrow) and cleaved PARP (magenta arrow) in U-118 control and EMP3 KO cells treated with 100 nM or 1 μM STS for 4 hours. **H** Concentration- response curve of U-118 control, EMP3 KO, and EMP3 KO + EMP3 cells treated with the EGFR inhibitor AZD9291 for 24 hours. Points represent mean percentage changes in CellTiter- Glo luminescence levels of treated cells relative to untreated cells, while error bars represent S.D. (n=3). **I** Caspase 3/7 levels in U-118 control, EMP3 KO, and EMP3 KO + EMP3 cells treated with 2.5 μM AZD9291 for 24 hours. Bars represent mean fold-changes in Caspase 3/7 luminescence readings of treated cells versus untreated controls. Error bars and dots represent S.D. and individual measurements, respectively (Welch’s ANOVA with Dunnett’s T3 multiple comparisons test; *P < 0.05).

Our -omics data indicated that several pathways and kinases related to EGFR signaling and tyrosine kinase inhibitor resistance are inhibited upon EMP3 depletion. To test whether this translates to increased sensitivity to kinase inhibition, we measured apoptotic rates in EMP3 KO and control cells upon pan-kinase inhibition with staurosporine (STS). STS is a broad-spectrum kinase inhibitor that binds to most kinases at submicromolar affinity [15]. Supporting our hypothesis, we observed that EMP3 depletion sensitizes U-118 and DK-MG cells to STS-induced apoptosis, as measured by higher caspase 3/7 activity (Fig. 6E and 6F) and greater cleaved PARP levels (Fig. 6G) in EMP3 KO cells. Thus, EMP3 may support the pro-survival activity of EGFR-dependent kinases, thereby allowing GBM cells to evade apoptotic cell death.

To further demonstrate the potential therapeutic relevance of EMP3 depletion, we also investigated whether EMP3 KO and targeted EGFR inhibition can have a synergistic effect in GBM. To test this, we measured cell viability and active caspase 3/7 levels in U-118 cells after treatment with AZD9291 (osimertinib). AZD9291 is a third-generation EGFR inhibitor that has shown efficacy against non-small-cell lung cancer [24]. Because it is highly brain-penetrant and can irreversibly inhibit both wild-type and mutant forms of EGFR, AZD9291 has been explored as a potential drug treatment for GBM [6, 9, 36]. Treatment of U-118 cells with AZD9291 for 24 hours reduced cell viability in a dose-dependent manner (Fig. 6H). The concentration-response curve exhibited a leftward shift upon EMP3 depletion, indicating a synergistic effect between EMP3 KO and EGFR-specific inhibition. This effect was reversed when EMP3 was re-expressed in EMP3 KO cells (Supplementary Fig. S7A), demonstrating that increased sensitivity to AZD9291 is a specific effect of EMP3 depletion. This effect was also reflected in the half-maximal inhibitory concentration (IC_50_) values, which was lower in EMP3 KO cells (IC_50_ = 3.65 µM) compared to control (IC_50_ = 4.37 µM) and EMP3 KO cells with the rescue construct (IC_50_ = 4.72 µM). Treatment with a sub-IC_50_ concentration of 2.5 µM AZD9291 for 24 hours also induced higher caspase 3/7 activity in EMP3 KO cells compared to control or EMP3 KOs with the rescue construct (Fig. 6I). Therefore, EMP3 contributes to therapeutic resistance against EGFR inhibition; conversely, targeting EMP3 may improve the effect of targeted EGFR inhibitors against GBM cells.

### EMP3 silencing in EGFR-high patient-derived glioblastoma stem cells increases susceptibility to CDK2 inhibition

GBMs are known to harbor both differentiated and stem like-cells [13, 57]. Our DK-MG and U-118 cultures closely mimic differentiated GBM cells, as these cell lines are grown in differentiation-promoting, serum-containing medium and exhibit morphological features akin to astrocytes. To further test whether our findings are applicable to stem-like GBM cells, we also generated patient-derived glioblastoma stem cell (GSC) models cultured as three-dimensional (3D) spheroids in serum-free stem cell conditions. We obtained EGFR-high (NCH1425) and EGFR-low (NCH644) GSCs (Supplementary Fig. S7B) and lentivirally transduced them with doxycycline-inducible non-targeting and EMP3-targeting short hairpin RNAs (shRNAs). Western blots confirm efficient knockdown of EMP3 in doxycycline-treated NCH644 and NCH1425 cells transduced with EMP3 shRNAs (Fig. 7A-B). We then proceeded to test whether loss of EMP3 synergizes with CDK2 inhibition in an EGFR-dependent manner. We hypothesized that if EMP3 facilitates CDK2 activity primarily through EGFR, then EMP3 knockdown should synergize with CDK2 inhibition only in EGFR-high GSCs. Conversely, EGFR-low GSCs expressing EMP3 shRNAs should not exhibit increased susceptibility to CDK2 inhibition, because the EMP3/EGFR/CDK2 signaling axis is non-existent in this context. Indeed, treatment with the selective CDK2 inhibitor K03861 synergized with EMP3 knockdown in a dose-dependent manner in NCH1425, but not in NCH644 GSCs (Fig. 7C-D). Thus, like in differentiated GBM cells, EMP3 also facilitates CDK2 activity in an EGFR-dependent manner in stem-like GSCs.

**Figure 7.**
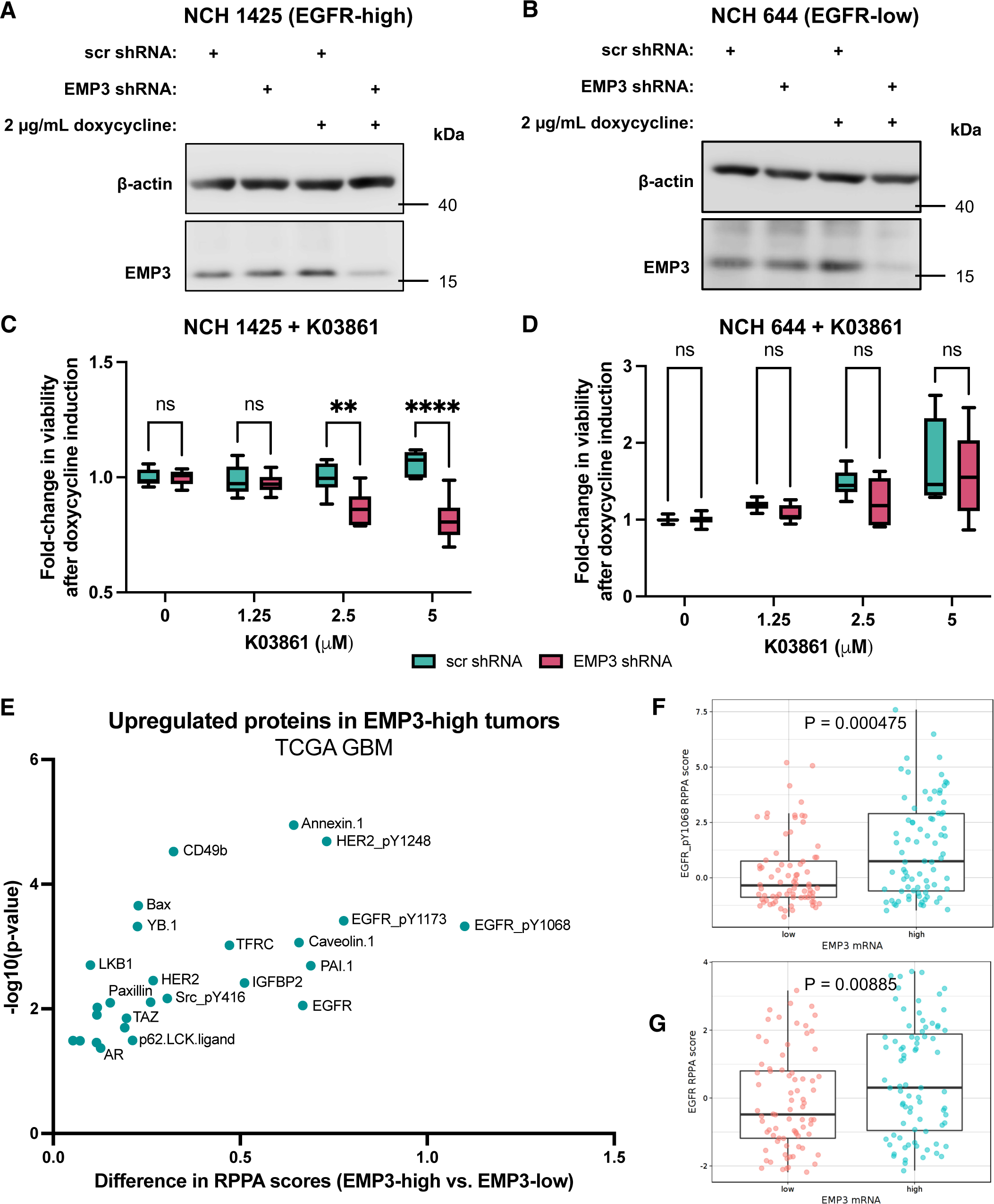
Validation of EMP3’s effects on EGFR and CDK2 using glioblastoma stem cells and TCGA data. **A, B** Western blots verifying successful EMP3 silencing in doxycycline-treated NCH1425 (A) and NCH644 (B) GSCs transduced with inducible EMP3 shRNAs. **C, D** Fold-change in the viability of NCH1425 (C) and NCH644 (D) GSCs after induction of shRNA expression and treatment with increasing concentrations of the CDK2 inhibitor K03861 (multiple Welch’s t-test; **P < 0.01, ****P < 0.0001). **E** Scatter plot showing upregulated proteins (difference in RPPA scores > 0; P < 0.05) in EMP3-high vs. EMP3-low TCGA IDH-wt GBM tumors. Tumors were classified into either group using the median EMP3 expression level based on the TCGA Agilent-4502A microarray data as cutoff. **F, G** Representative bar plots showing the RPPA scores of Tyr1068-phosphorylated (F) and total EGFR (G) in EMP3-high vs. EMP3-low IDH- wt GBMs. Figures were obtained from the GlioVis portal version 0.20 (http://gliovis.bioinfo.cnio.es/, accessed 09 July 2022).

### EMP3-high GBMs have elevated levels of total and phosphorylated EGFR

Lastly, to see whether our *in vitro* findings correspond to what is observed in actual tumor samples, we mined the TCGA GBM dataset using the GlioVis portal [4]. We searched for proteins upregulated in EMP3-high versus EMP3-low tumors (i.e., difference in reverse phase protein array (RRPA) scores > 0; P ≤ 0.05). Interestingly, both total and phosphorylated EGFR (Tyr1068 and Tyr1173) were among the top proteins with increased abundance in EMP3-high tumors (Fig. 7E). Other RTKs (e.g., HER2) and BioID2-identified EMP3 interactors (e.g., TFRC, CAV1) also had higher protein abundances in EMP3-high tumors, hinting at possible EMP3-dependent regulation of these proteins as well. The higher levels of Tyr1068- phosphorylated (Fig. 7F) and total EGFR (Fig. 7G) in EMP3-high IDH-wt GBMs are consistent with our *in vitro* findings demonstrating increased degradation of EGFR upon EMP3 depletion. In summary, EMP3-dependent maintenance of EGFR stability and activity is consistent across biochemical, transcriptomic, phosphoproteomic, and phenotypic levels and can be further correlated with clinical data from the TCGA.

## Discussion

In this study, we integrated protein-protein interaction (PPI) mapping with phosphoproteomics, transcriptomics, and functional characterization of CRISPR/Cas9 KO cells to examine the molecular function of EMP3 in IDH-wt GBM. Our results show that EMP3 stabilizes EGFR, a frequently overactivated oncogene in IDH-wt GBM. Consistent with previous studies that showed inhibition of RTK signaling in non-glioma cells upon EMP3 silencing [11, 21, 59], we observed higher EGF-induced EGFR degradation in our EMP3 KO cells. Building on these findings, we further elucidated a novel link between the stabilization of EGFR and EMP3’s trafficking function. Specifically, we have shown that EMP3 restricts EGFR trafficking into degradative endosomes, as loss of EMP3 promoted the association between ligand-activated EGFR and the late endosomal marker RAB7. These *in vitro* findings are consistent with TCGA data, which show reduced levels of both total and phosphorylated EGFR in GBMs with low EMP3 expression. In our experimental model, increased EGFR degradation was rescued by overexpression of the retromer component TBC1D5, a novel EMP3 interactor identified by our BioID2 screen. Other retromer components, including the sorting nexins SNX1 and SNX2, have been previously linked to the regulation of EGFR trafficking and stability [17, 34, 63]; however, to our knowledge, this is the first report of TBC1D5’s involvement in regulating EGFR dynamics. As a RAB7 GTPase-activating protein (GAP), TBC1D5 inhibits several RAB7- mediated processes, including RAB7 localization to late endosomes [26, 35, 48, 49]. Critically, RAB7 mediates EGFR shuttling from late endosomes to lysosomes, thereby facilitating receptor degradation [1, 14]. Given these past findings and our present results, we propose that EMP3 facilitates TBC1D5 recruitment into maturing endosomes, where the latter could inactivate RAB7 and thus restrict the progression of internalized EGFR cargoes towards lysosomal degradation (Fig. 8). This model is consistent with previous work demonstrating TBC1D5-mediated retrieval of other receptors ITGA5, ITGB1, and IGF2R away from endolysosomal degradation [25]. Interestingly, we also identified these receptors as EMP3 interactors in our BioID2 screen, indicating that the EMP3-TBC1D5 complex may stabilize a broader set of GBM receptors beyond EGFR. Additional work will be necessary to systematically identify what other receptors are regulated by the EMP3-TBC1D5 interaction.

**Figure 8.**
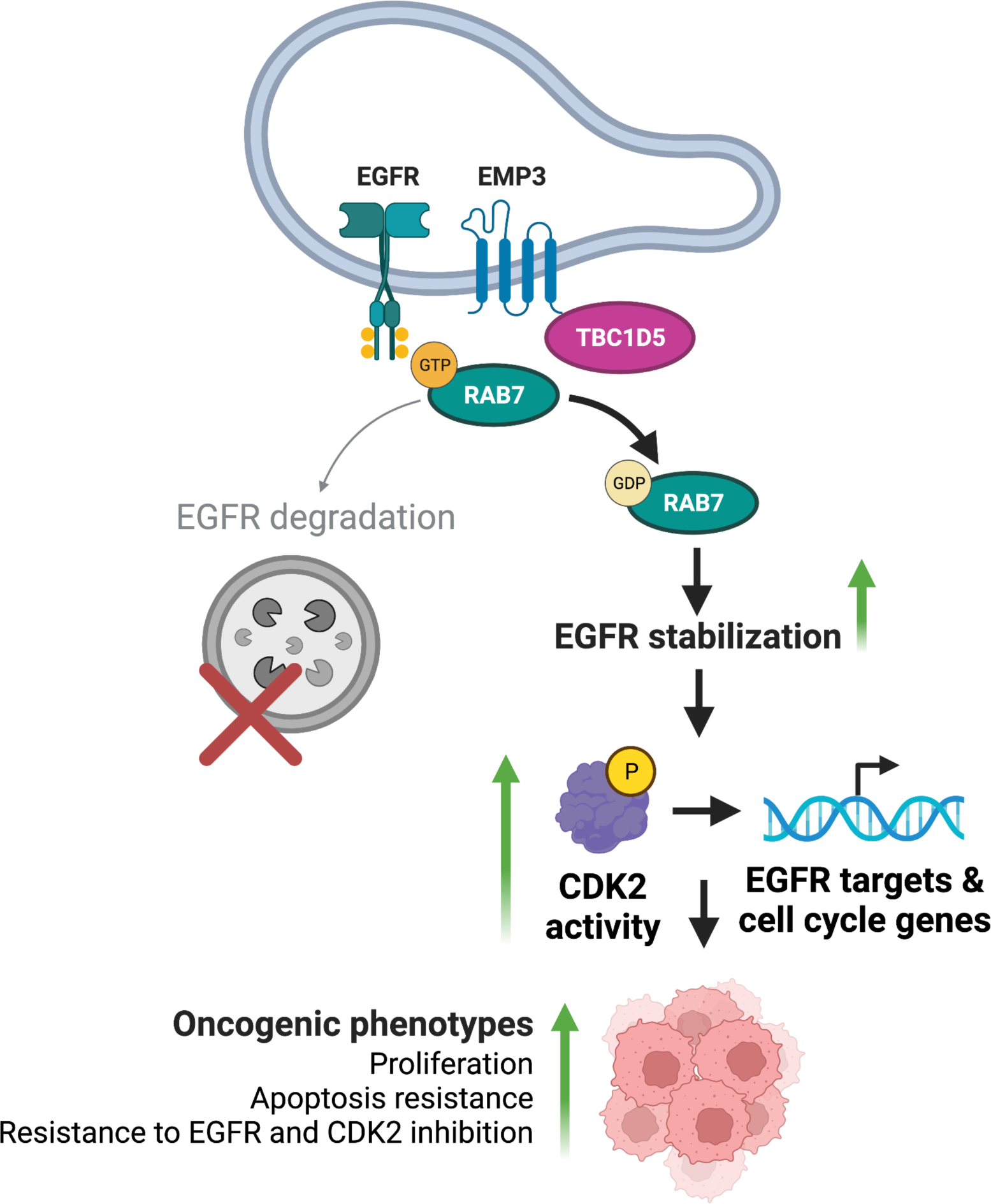
Proposed model of EMP3 function in GBM. EMP3 interacts with TBC1D5, and the resulting complex inactivates RAB7 in late endosomes. Inactive RAB7 is unable to facilitate EGFR degradation, and EGFR is stabilized. EMP3-dependent stabilization of EGFR sustains downstream signaling via the EGFR effector CDK2. This cascade culminates in the transcription of EGFR-responsive genes involved in cell cycle progression. Ultimately, these mechanisms ensure sustained proliferation, apoptosis resistance, and reduced susceptibility to targeted kinase inhibitors. Figure created with BioRender.

In addition to these mechanistic findings, our study also demonstrates how impaired EGFR trafficking and reduced EGFR stability upon EMP3 KO inhibit downstream oncogenic signaling. First, our phosphoproteomic data indicates that EMP3 KO leads to the inactivation of the cyclin-dependent kinase CDK2. CDK2 has been previously shown to be downstream EGFR target in glioblastoma cell lines [7], and its transient activation by AKT is known to be a critical step in the induction of cell cycle progression [38]. Apart from promoting *in vitro* proliferation of GBM cells, CDK2 also facilitates *in vivo* tumor growth as well as resistance to apoptosis induced by radiotherapy [58]. Consistent with concomitant EGFR and CDK2 inhibition, our gene expression data demonstrate that EMP3 KO leads to the downregulation of EGFR-responsive genes that are involved in DNA replication and cell cycle regulation. In line with these phosphoproteomic and transcriptomic changes, we show that EMP3 KO cells have reduced cellular proliferation and blunted mitogenic response to EGF. The mechanism and signaling alterations defined here are likely to be broadly applicable to other non-glioma tumor entities, as their RTK signaling outputs have also been shown to be supported by EMP3 [11, 21, 59].

Importantly, our results shed light on how EMP3 could modulate the outcome of targeted therapies. Our data indicate that loss of EMP3 impairs the transcription of EGF/EGFR-dependent genes that mediate tyrosine kinase inhibitor resistance. Consistent with this, EMP3- expressing cells display lower sensitivities to pan-kinase and EGFR-specific inhibition compared to EMP3 KO cells. Furthermore, the differential susceptibility of NCH1425 GSCs expressing scrambled and EMP3-targeting shRNAs to the CDK2-specific inhibitor K03861 demonstrates that EMP3 is sufficient to confer resistance against this drug as well. It is well known that GBM cells employ redundant mechanisms to ensure that oncogenic signaling is sustained even after aggressive treatment [22]. By sustaining EGFR/CDK2 activity and by being expressed at such high levels in this tumor entity, EMP3 provides an additional layer of resistance that protects tumor cells from targeted kinase inhibitors. This, along with other resistance mechanisms, could explain why kinase inhibitors have been largely unsuccessful in various clinical trials [43, 52]. It will be worthwhile for future GBM drug discovery screens to explore pharmacological agents inhibiting EMP3. While EMP3 is not a classical oncogene driver, its crucial role in supporting EGFR/CDK2 signaling makes it an appealing addition to traditional somatically altered targets.

Our BioID2-based proximity labeling approach also revealed several other EMP3-proximal proteins, including GBM-relevant membrane receptors (e.g., CD44, integrins, SLCs), signaling adaptors, and other trafficking regulators. This rich dataset provides a wealth of testable hypotheses for future mechanistic investigations on both tumor-intrinsic and tumor-extrinsic effects of EMP3. Given EMP3’s emerging role in the tumor immune microenvironment [10] and its high expression in both GBM and tumor-infiltrating macrophages and T cells [42], it would be interesting to investigate how its membrane interactions and/or receptor trafficking function could ultimately influence cancer cell-immune cell crosstalk in GBM. Also of interest is the potential distinct role of glycosylated EMP3, which appears to preferentially associate with inner mitochondrial membrane proteins based on our BioID2 data. Previous studies have shed light on glycan-dependent localization and function of glycosylated mitochondrial proteins [5, 33]. Considering the role of glycosylation in cancer [44], it will be important to investigate the potential role of glycosylated EMP3 in the mitochondria and how it could influence GBM independent of EMP3’s effects on EGFR.

In conclusion, this study identifies novel interacting partners of EMP3, thereby highlighting its multi-localizing nature while clarifying the subcellular context in which it could operate as a tumor-promoting protein. Specifically, we unravel a novel EMP3-dependent trafficking mechanism that maintains EGFR activity. By associating with the RAB7 GAP TBC1D5, EMP3 restricts the late endosomal trafficking and inhibits the degradation of activated EGFR receptors. Such a mechanism ensures the maintenance of the EGFR/CDK2 signaling axis, which promotes tumor cell proliferation and provides IDH-wt GBM tumor cells with an additional layer of resistance against targeted EGFR/CDK2 inhibition.

## Declarations

### Ethics approval and consent to participate

No consent to participate was required for this study.

### Consent for publication

All authors consented to publication.

### Availability of data and materials

The data generated in this study are available within the article and its supplementary files.

### Author’s Contributions

**AAM**: Conceptualization; Formal analysis; Investigation; Methodology; Software; Validation; Visualization; Writing – original draft. **AK**: Investigation; Methodology; Validation; Formal analysis. **NB**: Investigation; Methodology; Formal analysis. **LD**: Validation. **AC**: Methodology; Resources. **DK**: Methodology; Software. **MS**: Data curation; Methodology; Software. **DH**: Data curation; Methodology; Project administration. **RW**: Methodology. **CH**: Funding acquisition; Supervision. **CH-M**: Resources; **AvD**: Funding acquisition; Supervision; Writing – review & editing. **SP**: Conceptualization; Funding acquisition; Project administration; Resources; Supervision; Writing – review & editing.

## Acknowledgements

We thank Susanne Gärtner for the technical assistance and Dr. Julia Zaman, Dr. Peter Angel, and Dr. Francesca Ciccolini for providing valuable feedback on this study. We also thank David Vonhören for his assistance in the Perseus analysis of the phosphoproteomics data. We also thank the DKFZ Light Microscopy Facility and the following units of the DKFZ Genomics and Proteomics Core Facility for their critical support to this study: Antibody Unit, Cellular Tools/Vector and Clone Repository Unit, MS-based Protein Analysis Unit, and Microarray Unit. Figures 8 and Supplementary Figures S1 and S4A were created with BioRender.

## Supplementary Figure Legends

**Supplementary Figure S1.**
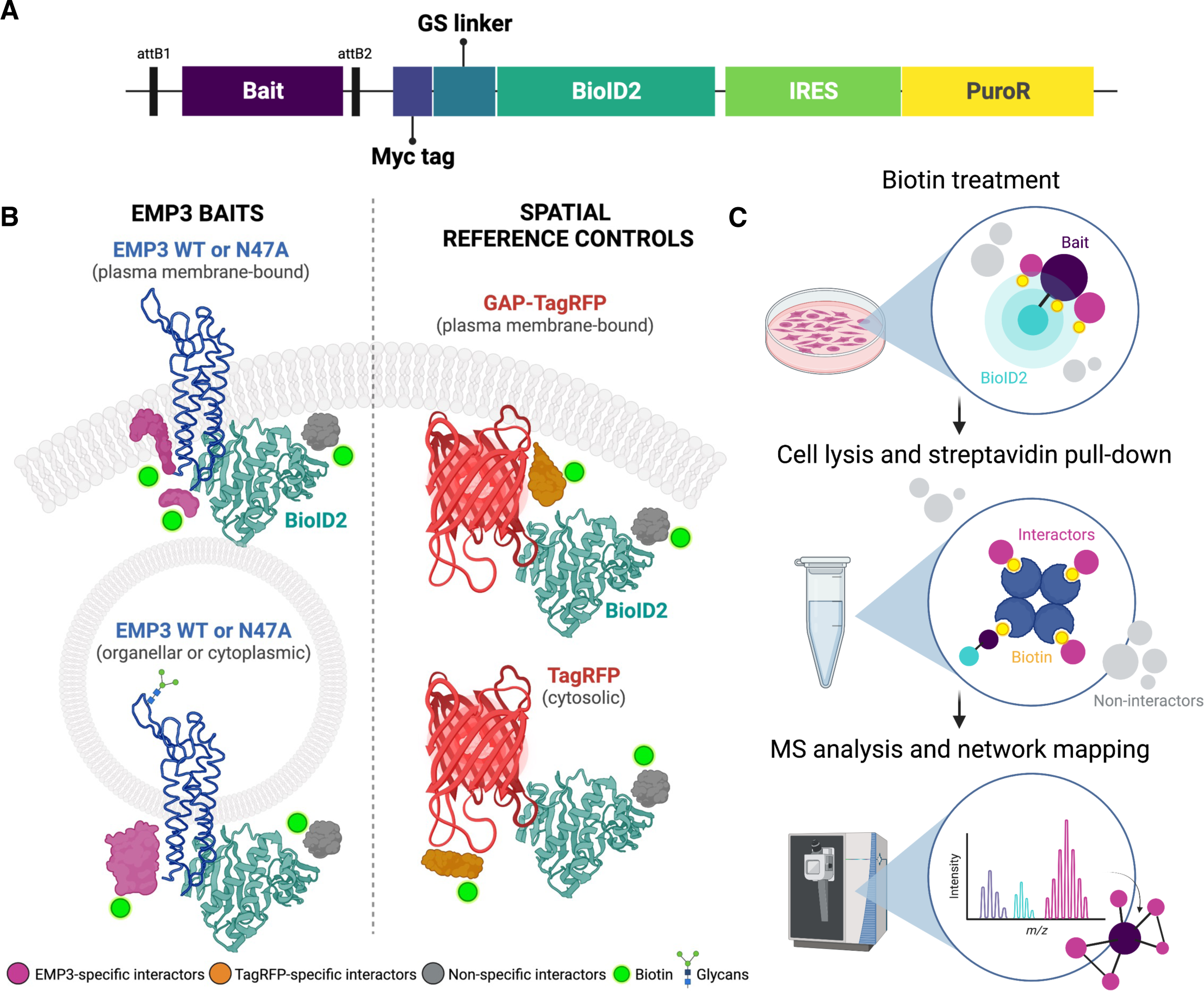
**A** Schematic diagram of the Gateway construct expressing the bait protein fused to a C-terminal Myc glycine-serine (GS) linker-BioID2 tag. An IRES-PuroR cassette follows the coding sequence of the fusion protein. **B** Experimental design showing baits used in the BioID2 experiment. Wild-type and N47A mutant versions of EMP3 tagged with Myc-Linker-BioID2 at the C-terminal end were used as experimental baits. EMP3 baits may localize in the plasma membrane or within unidentified cytoplasmic compartments. TagRFP and GAP-TagRFP fusion proteins, which localize in cytoplasm and the plasma membrane, respectively, were used as spatial reference controls. **C** Workflow detailing experimental steps from biotin treatment to mass spectrometry (MS) analysis of identified EMP3-proximal proteins. U-118 cells stably transfected with BioID2 constructs were treated with 50 μM biotin for 18 hours to induce BioID2-mediated biotinylation. Biotinylated proteins were purified by streptavidin pull- downs, and the eluates were subjected to MS and network analysis. Figures created with BioRender.

**Supplementary Figure S2.**
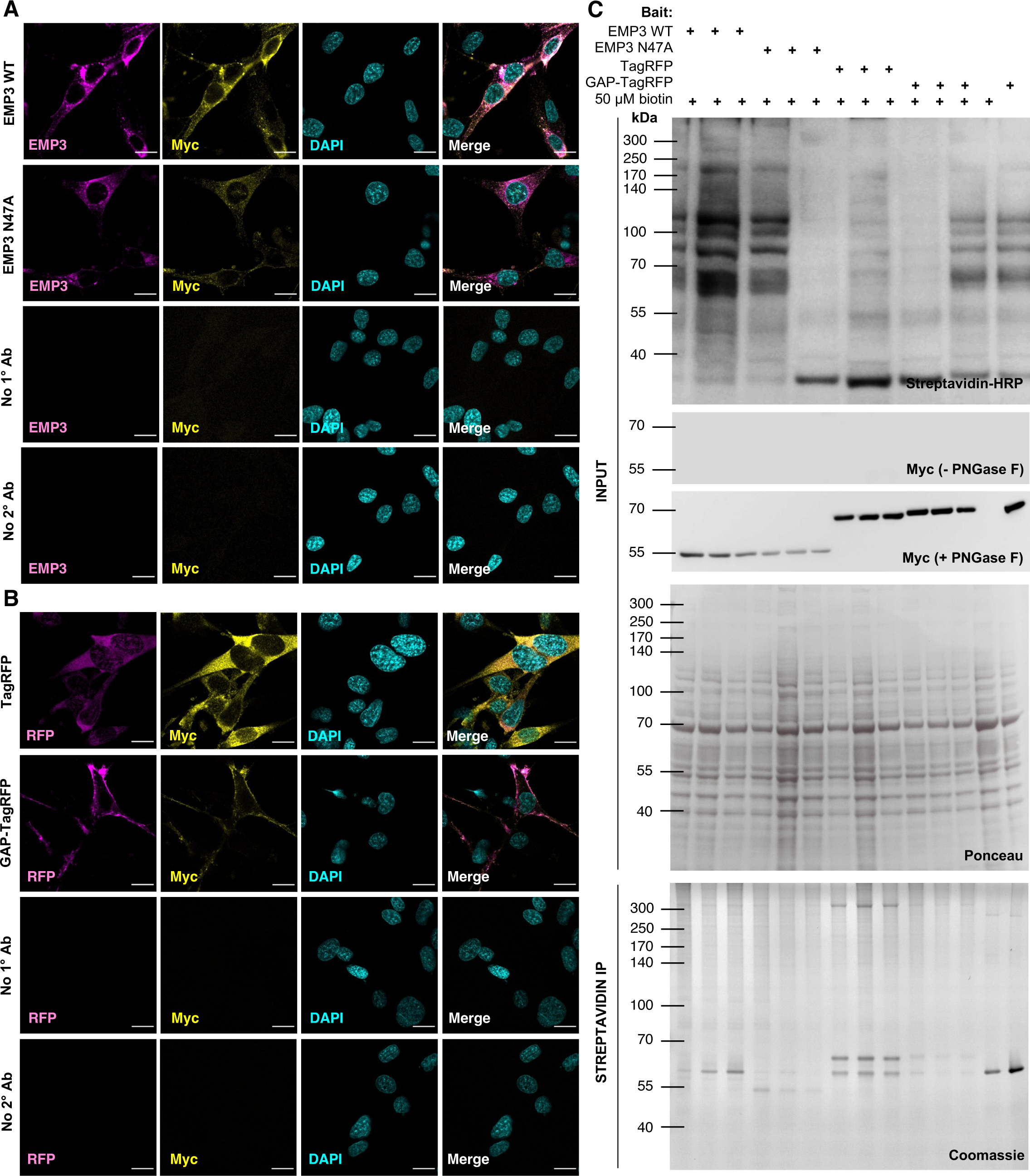
**A** Immunofluorescence (IF) staining of Myc-Linker-BioID2-tagged EMP3 WT and N47A proteins in U-118 cells. Antibodies against EMP3 (magenta) and Myc (yellow) were used to stain BioID2 fusion proteins. DAPI (cyan) was used for nuclear staining. Scale bar = 20 μm. **B** IF staining of Myc-Linker-BioID2-tagged TagRFP and GAP-TagRFP proteins in U-118 cells. RFP fluorescence (magenta) was visualized alongside Myc (yellow) and nuclear staining (DAPI). U-118 cells without either TagRFP constructs were used as controls. Scale bar = 20 μm. **C** Validation of BioID2-based proximity labeling. Bait expression was confirmed by immunoblotting for the Myc tag. Biotinylation of bait-proximal proteins was verified by streptavidin-HRP blots of the input lysate. Coomassie staining was performed on the eluates to confirm purification of biotinylated proteins.

**Supplementary Figure S3.**
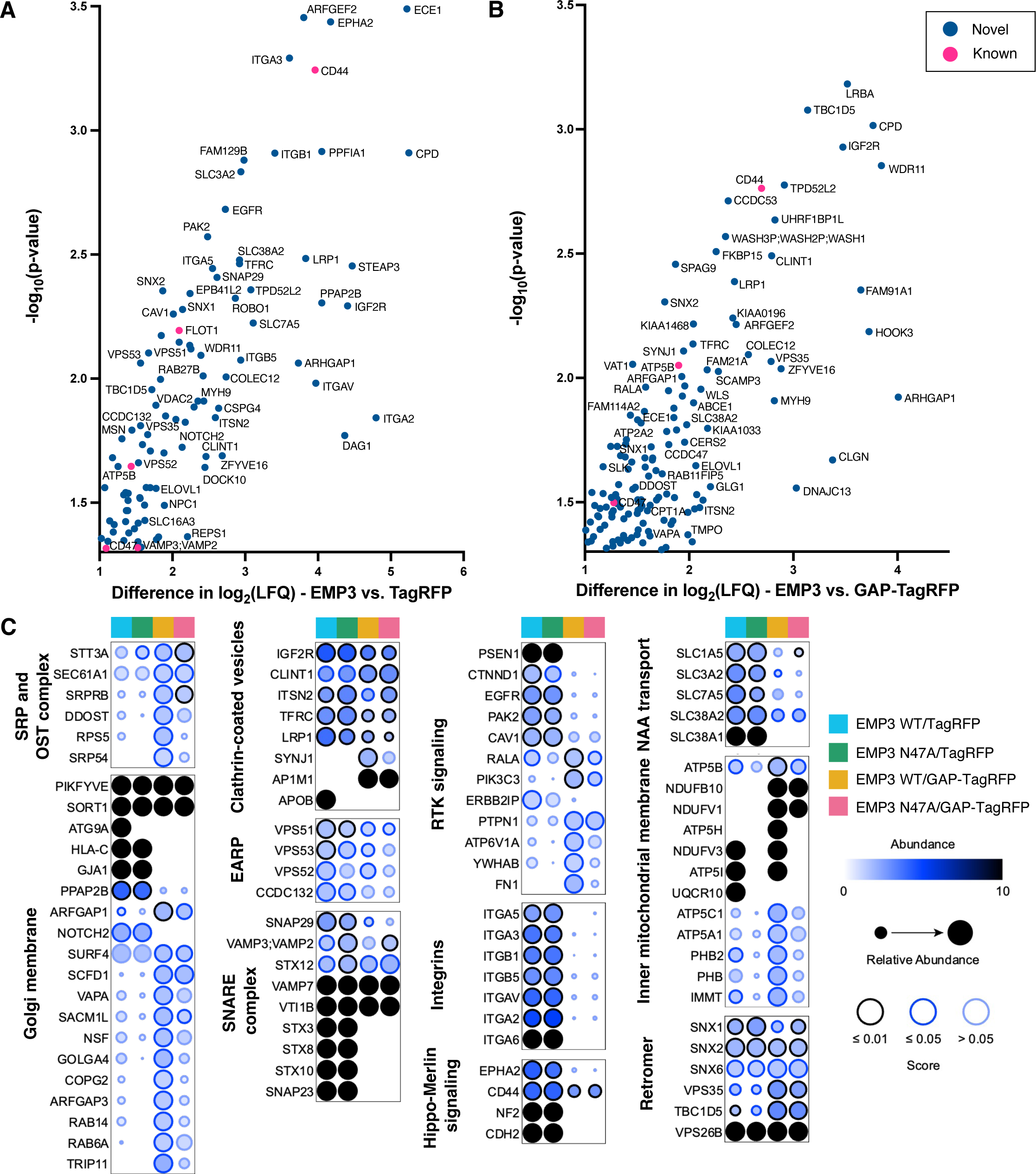
**A, B** Scatter plots showing EMP3-proximal proteins (i.e. difference in log2(LFQ) ≥ 1, Welch’s t-test p-value ≤ 0.05) identified using EMP3 WT as bait and TagRFP (A) or GAP-TagRFP (B) as the spatial reference control. Blue dots indicate novel EMP3-interacting partners, while pink dots represent previously identified EMP3 interactors based on the EMP3 yeast two-hybrid screen by Christians et al. and the BioGRID database (v4.4). **C** Dot plots showing relative enrichment of EMP3-proximal proteins, filtered and grouped in Cytoscape as previously described, across possible pairwise comparisons. Dot colors and sizes correspond to the difference in log2(LFQ) values of EMP3 baits versus spatial reference controls. The selected abundance cap automatically assigns a black fill color to proteins uniquely purified in EMP3 WT and N47A pull-downs while allowing visualization of the range of possible enrichment values. Border colors correspond to significance levels. All dot plots were generated using ProHits-viz.

**Supplementary Figure S4.**
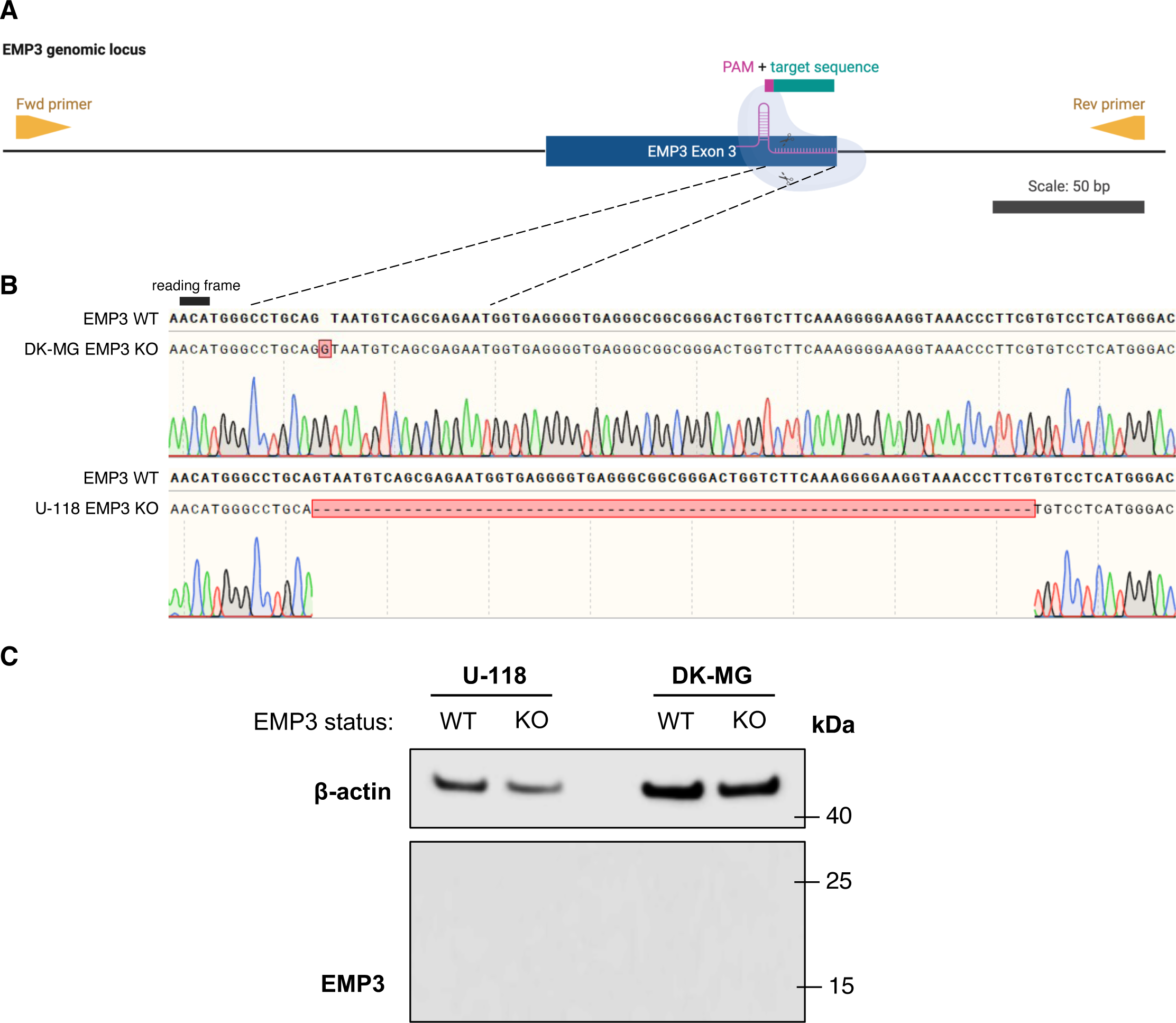
**A** Schematic diagram showing the EMP3 genomic locus containing Exon 3 flanked by intronic sequences on either side, and the relative positions of the guide RNA (gRNA) target sequence, PAM site, and primer-binding sites for forward (Fwd) and reverse (Rev) sequencing primers. Scale bar = 50 bp. Figure created with BioRender. **B** Sequencing results confirm the insertion of a single guanine nucleotide 2 bp downstream of the putative gRNA cut site in the DK-MG EMP3 KO cell line and a 71-bp deletion after the cut site in U-118 EMP3 KO cells. Both alterations should induce nonsense-mediated mRNA decay of the EMP3 transcript. **C** Western blots confirming proper EMP3 KO at the protein level in both cell lines.

**Supplementary Figure S5.**
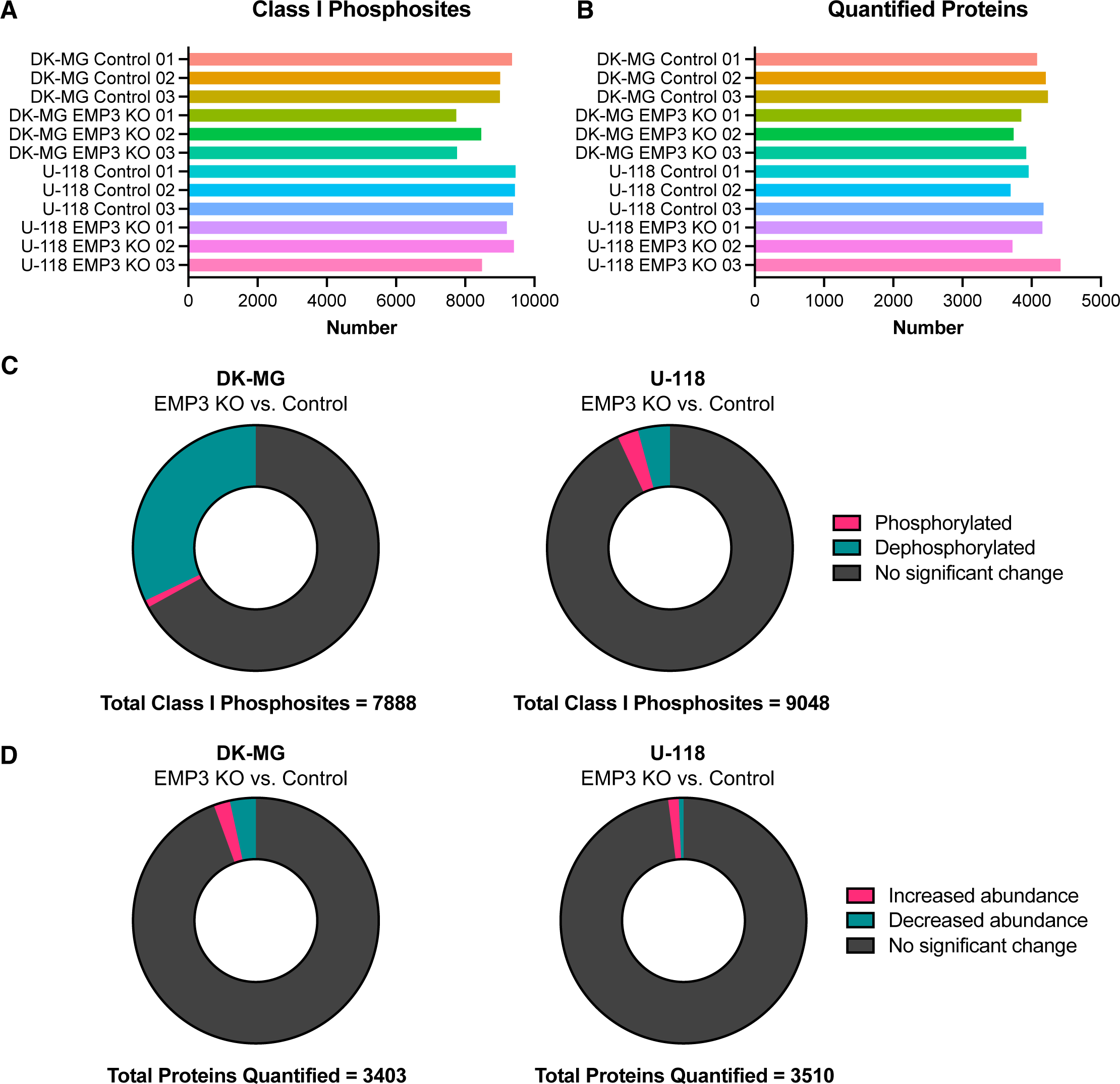
**A, B** Bar plots showing the total number of class I phosphosites (i.e., localization probability ≥ 0.75) identified (A) and total number of proteins quantified in each sample (B) after mass spectrometry analysis. **C** Donut charts showing the distribution of phosphorylation changes occurring in class I phosphosites that were detected in at least 2 out of 3 replicates of each condition (i.e., control or EMP3 KO) in DK-MG (left) and U-118 (right) cells. Phosphorylated - log_2_-FC ≥ 1, FDR p-value ≤ 0.05; Dephosphorylated - log_2_-FC ≤ -1, FDR p-value ≤ 0.05; No significant change – [log_2_-FC| < 1 and/or FDR p-value > 0.05. **D** Donut charts showing the distribution of proteins that were quantified in at least 2 out of 3 replicates of each condition (i.e., control or EMP3 KO) in DK-MG (left) and U-118 (right) cells. Increased abundance – log_2_- FC ≥ 1, FDR p-value ≤ 0.05; Decreased abundance – log_2_-FC ≤ -1, FDR p-value ≤ 0.05; No significant change – [log_2_-FC| < 1 and/or FDR p-value > 0.05.

**Supplementary Figure S6.**
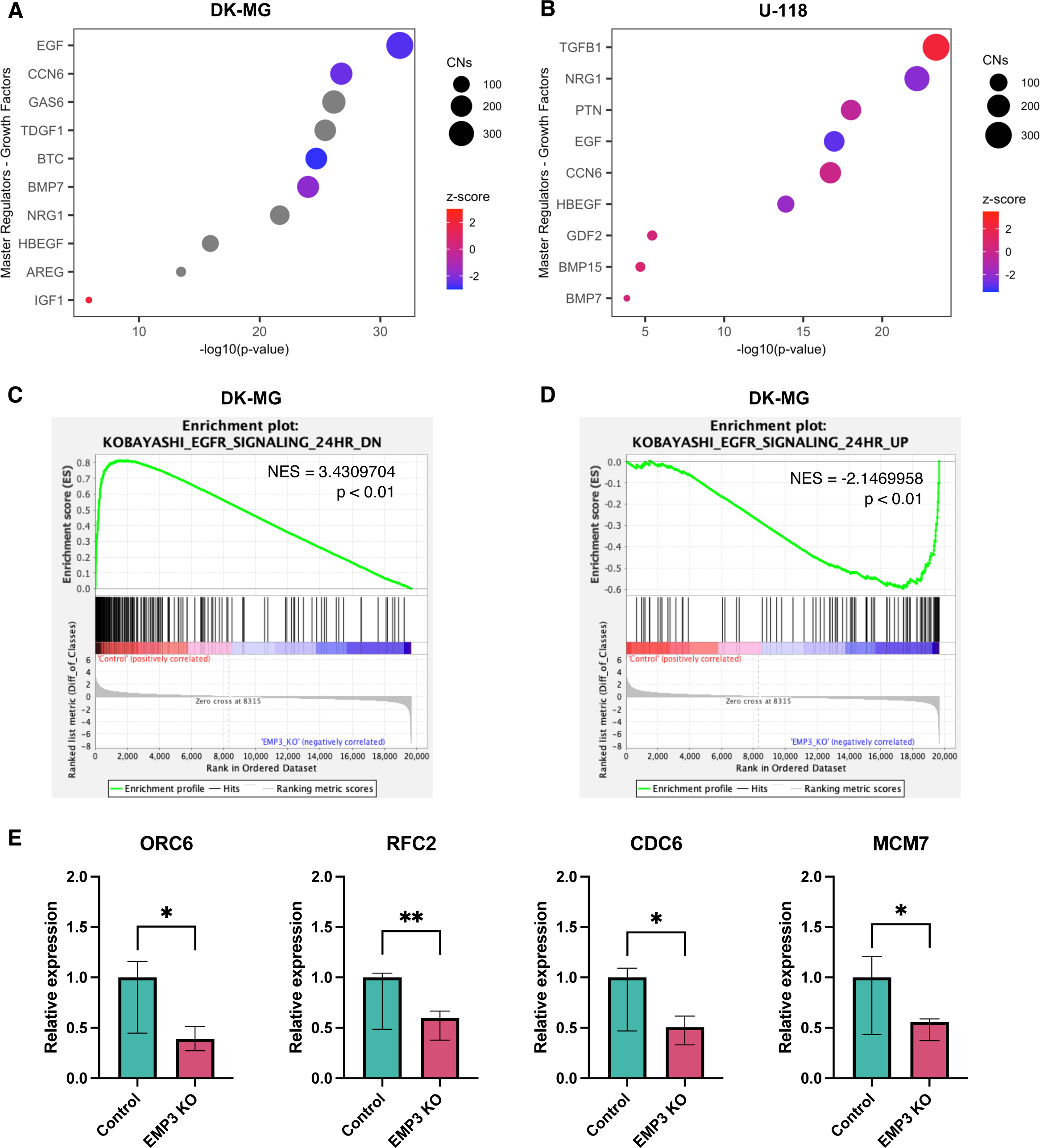
**A, B** Growth factors identified to be significantly enriched master regulators (MRs) based on DEGs in DK-MG (A) and U-118 (B) EMP3 KO relative to controls. MRs putatively regulating the input genes are listed on the y-axis and ordered according to significance. Circle sizes represent the number of associated causal networks (CNs) per MR, while the color scale indicates the activation z-score of each MR (red – active; blue– inactive). **C, D** GSEA analysis showing upregulation of the KOBAYASHI_EGFR_SIGNALING_24HR_DN (C) and KOBAYASHI_EGFR_SIGNALING_24HR_UP (D) gene sets in DK-MG control and EMP3 KO cells, respectively. Genes were sorted from left to right based on the difference of the log_2_ expression levels between control and EMP3 KO cells. Vertical black bars indicate the location of the genes contributing to the enrichment scores (ES). The ES, which indicate upregulation (ES > 0) or downregulation (ES < 0) of a certain gene, are plotted on the y-axis. NES: normalized enrichment score. **E**, qPCR results validating the downregulation of selected genes in DK-MG EMP3 KOs (unpaired one-tailed t-test; * P = < 0.05; ** P = < 0.01).

**Supplementary Figure S7.**
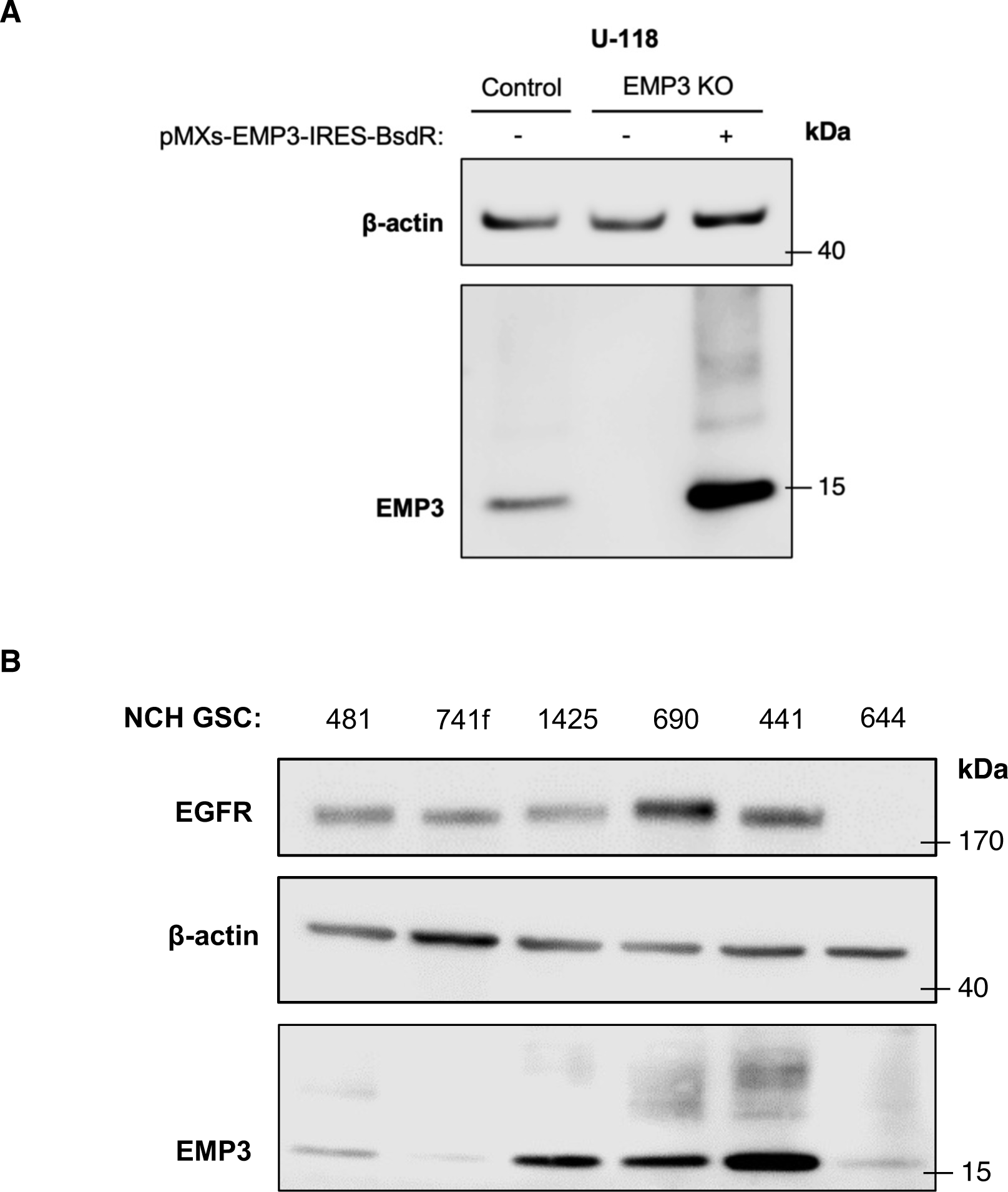
**A** Western blot confirming proper re-expression of EMP3 in U-118 EMP3 KO cells. EMP3 KO cells were stably transfected with pMXs-EMP3-IRES-BsdR to restore EMP3 expression. **B** Western blot showing EMP3 and EGFR levels in a panel of six patient-derived GSC lines. NCH1425 (EGFR-high) and NCH644 (EGFR-low) were further used in the study.

