## Supplementary Information for "EMP3 sustains oncogenic EGFR/CDK2 signaling by restricting receptor degradation in glioblastoma"

**TABLE OF CONTENTS**


| Page Number | Table | Table Title |
|---|---|---|
| 2 | Table 1 | Primers used in this study |
| 3 | Table 2 | List of antibodies used in this study |

Table 1. Primers used in this study

| Product | Forward Primer (5‘ to 3‘) | Reverse Primer (5‘ to 3‘) |
|---|---|---|
| pMXs-Gateway | CTACGGCTACACTAGAAGAACAGTATTTGGTATC | CCCGTCAACCACTTTGTACAAGAAAGCTG |
| Myc-Linker | TGTACAAAGTGGTTGACGGGGAACAAAAACTCATCTCAGAAGAGGATCTCGACGGTGGAGGCGGGTCTGGA | GGTTCTTGAAACCGGTCGATCCACCGCC |
| Linker-BioID2 | ATCGACCGGTTTCAAGAACCTGATCTGG | AATTTACGTAGCGGCCGCGGTTAGCTTCTTCTCAGGCTG |
| IRES-PuroR | CCGCGGCCGCTACGTAAATT | TTCTTCTAGTGTAGCCGTAGTTAGGCCACC |
| GAP-TagRFP in pDONR201 | CTGTGCTGTATGAGAAGAACCAAACAGAGCGAGCTGATTAAGGAGAACATGC | GTTCTTCTCATACAGCACAGCATGGTGGAGCCTGCTTTTTTGT |
| TBC1D5 R169A | GTCAAAGCAACGTTTCCTGAAATGCAGTTTTTCCA | GGAAACGTTGCTTTGACATCTTGTTCAATCATTGATCGAAGT |
| TBC1D5 Q204A | TTATAAAGCGGGCATGCACGAACTGTTAGC | TGCATGCCCGCTTTATAAAGCAACTGCTCGTTTTCTCTGG |
| EMP3 shRNA | GTGAAGCCACAGATGTATAGAGCTGGAACATGAACAGGAAGTGCCTACTGCCTCGGAATTCTAAAGTAG | TACATCTGTGGCTTCACTATAGAGCTGGAACATGAACAGGAATCGCTCACTGTCAACAGCAATATAC |

Table 2. List of antibodies used in this study

| Antibody Target | Host | Company | Catalog No. | Dilution |
|---|---|---|---|---|
| Phospho-EGF receptor (Tyr1068) | rabbit | Cell Signaling Technology (CST) | 3777 | WB: 1:1000 |
| Phospho-EGF receptor (Tyr1068) | mouse | Thermo Fisher | MA5-15199 | PLA: 1:100 |
| EGFR | rabbit | CST | 4267 | WB: 1:1000<br>PLA: 1:50 |
| PARP | rabbit | CST | 9542 | WB: 1:1000 |
| β-actin | rabbit | CST | 4970 | WB: 1:1000 |
| Myc | rabbit | CST | 2278 | WB: 1:1000<br>IF: 1:200 |
| FLAG | mouse | Sigma-Aldrich | F1804 | WB: 1:1000<br>IF: 1:500 |
| EMP3 | mouse | DKFZ Antibody Core Facility | K158/2 | WB, IF, and PLA: undiluted |
| EMP3 | mouse | DKFZ Antibody Core Facility | K158/8 | IF: 1:10 |
| RAB7 | rabbit | CST | 9367 | PLA: 1:100 |
| CLINT1 | rabbit | Thermo Fisher | PA5-60308 | PLA: 1:50 |
| SNX1 | rabbit | Thermo Fisher | MA5-34808 | PLA: 1:100 |
| SNX2 | rabbit | Thermo Fisher | PA5-83367 | PLA: 1:100 |
| TBC1D5 | rabbit | Abcam | ab203896 | PLA: 1:200 |
| VPS53 | rabbit | Thermo Fisher | PA5-55079 | PLA: 1:50 |
| SOX2 | rabbit | CST | 23064 | PLA: 1:400 |
| CDK2 | rabbit | CST | 18048 | WB 1:1000 |
| Rabbit IgG | goat | CST | 7074 | WB: 1:4000 |
| Mouse IgG | horse | CST | 7076 | WB: 1:4000 |
