## Supplementary Materials and Methods for "EMP3 sustains oncogenic EGFR/CDK2 signaling by restricting receptor degradation in glioblastoma"

**Plasmid cloning**

*Generation of BioID2 plasmids*

The BioID2 destination vector was generated by NEBuilder® HiFi DNA Assembly. Briefly, the following DNA fragments were amplified by PCR using the primers listed in Supplementary Table 1: 1) pMXs-Gateway backbone, 2) overlapping Myc-Linker and Linker-BioID2 cassettes, 3) IRES-PuroR resistance marker. The fragments corresponding to Myc-Linker-BioID2 were amplified from a BioID2 template plasmid that was a kind gift from Dr. David Reuss. The following primers were used to amplify the Myc-Linker and Linker-BioID2 fragments: Meanwhile, the pMXs-Gateway backbone and IRES-PuroR resistance marker were amplified from a pMXs-GW-IRES-PuroR plasmid provided by the DKFZ Vector and Clone Repository.

To assemble the four overlapping DNA fragments, 0.125 pmol of each PCR product were pooled together, mixed with an equivalent volume of NEBuilder® HiFi DNA Assembly Master Mix (New England Biolabs E2621S), and incubated at 50°C for 1 hour. To digest the PCR templates, the reaction was further treated with DpnI (Invitrogen IVGN0106) for 30 minutes at 37°C prior to heat-inactivation at 80°C. The reaction was then transformed into ccdB survival competent cells (Invitrogen A10460) and plasmids were extracted using standard plasmid preparation procedures. Plasmid sequencing confirmed proper insertion of the Myc-Linker-BioID2 cassette between the pMXs-Gateway and IRES-PuroR segments, thereby generating the pMxs-GW-Myc-Linker-BioID2 destination vector. To generate the bait constructs, Gateway LR reactions were performed using the new BioID2 destination vector and existing pDONR plasmids harboring the EMP3 and TagRFP coding sequences in our lab. The GAP-TagRFP bait construct was likewise generated by LR reaction after introduction of the GAP-43 membrane-targeting sequence (MLCCMRRTKQ) into the N-terminal end of the TagRFP coding sequence by site-directed mutagenesis.

*Generation of EMP3 and TBC1D5 rescue constructs*

A TBC1D5 Gateway entry clone in pDONR223 was obtained from the Vector and Clone Repository of the DKFZ Cellular Tools Core Facility. The R169A/Q204A mutant was then generated by inverse PCR-based site-directed mutagenesis using the primers specified in Supplementary Information Table 1. Both wild-type and mutant TBC1D5 coding sequences were then subcloned in pMXs-GW-FLAG-IRES-

BsdR using a standard Gateway LR reaction. Similarly, the EMP3 coding sequence in pDONR221 was subcloned into the same destination vector.

*Generation of EMP3 shRNA plasmid*

To design the EMP3 shRNA construct, the top microRNA-adapted shRNA (shRNA-miR) cassette targeting EMP3 (target sequence: TCCTGTTCATGTTCCAGCTCTA) was first selected using miR_Scan [4]. The EMP3-targeting shRNA construct was then cloned via site-directed mutagenesis of a non-targeting scrambled shRNA in a pENTR1A backbone. using the primers listed in Supplementary Information Table 1. The shRNA cassette was then digested using Anza MluI (Thermo Fisher Scientific IVGN0286) and XhoI (Thermo Fisher Scientific IVGN0086) restriction enzymes and subcloned into a MluI/XhoI double-digested and alkaline phosphatase-treated (Thermo Fisher Scientific IVGN2208) pTRIPZ backbone (Horizon Discovery, Cambridge, UK) using the Anza T4 DNA ligase master mix (Thermo Fisher Scientific IVGN2104). Resulting constructs were sequence-verified by Sanger sequencing (Azenta Life Sciences)

**Generation of CRISPR/Cas9-based EMP3 knockout cell lines**

CRIPSR/Cas9 vectors pCas9-PuroR/EMP3 were constructed by inserting the guide RNA template sequence targeting EMP3 exon 3 (Forward Primer: CACCGATTCTCGCTGACATTACTGC, Reverse Primer: AAACGCAGTAATGTCAGCGAGAATC) into pSpCas9(BB)-2A-Puro (PX459) V2.0 (a kind gift from Feng Zhang; Addgene plasmid ID #62988) following a previously published protocol [5]. U-118 and DK-MG cells were transfected with pCas9-Puro/EMP3 using Lipofectamine 3000 (Thermo Fisher Scientific L3000001). After 24 hours, cells were briefly put under selection pressure with 1 µg/mL puromycin for 48 hours. After this period, the medium was changed back to normal growth medium without puromycin. Surviving single cell clones were marked and allowed to grow into colonies. Colonies potentially containing more than one clone were picked, diluted, and re-seeded to obtain single-cell clone colonies. Single-cell clone colonies were then picked and expanded. After sufficient cell numbers were reached, cells were detached with TrypLE Select (Thermo Fisher Scientific 12563011) and re-seeded into 24-well plates. EMP3 and Cas9-FLAG protein expression was determined by immunofluorescence using the EMP3 158/8 (DKFZ Antibody Core Facility) and anti-FLAG M2 antibodies (Sigma-Aldrich F1804), respectively. Only cell clones without visible expression of both EMP3 and Cas9-FLAG were selected for following experiments. After further expansion, genomic DNA from the selected candidate cell lines was extracted. The CRISPR/Cas9-targeted region was amplified by PCR (Forward Primer: TTAGCTCTACCTCCGATGCC, Reverse Primer: CGCCCACTCCAACTTTGTT) and TA-cloned into pGEM-T Easy plasmid vector (Promega). Ligation reactions were then transformed into *E.coli* DH5α and plated onto LB agar plates. After colony formation, colonies were picked and plasmids were extracted. The vector inserts were checked for indels by Sanger sequencing (GATC, Ebersberg, Germany).

**Stable plasmid transfection**

Transfection of GBM cells were performed using X-tremeGENE 9 DNA transfection reagent (Roche XTG9-RO). For each transfection, $2.0 \times 10^5$ cells were seeded in 2 mL of DMEM maintenance medium in one well of a 6-well plate. Once the cells are 70-80% confluent, cells were transfected by the addition of the transfection mix. Briefly, 2000 ng of plasmid DNA and 6 µL X-tremeGENE 9 were added to 200 µL of Opti-MEM Reduced Serum Medium (Gibco 31985-062) and the resulting mix was incubated at room temperature for 15 minutes. Cells were then transfected by the addition of the mix and then incubated at 37°C, 5% $CO_2$ for 72 hours to allow sufficient plasmid expression. Stably transfected cells were selected by continuous treatment of the cells with the appropriate antibiotic-containing medium. Selected cells were then characterized by Western blotting and maintained in cell culture flasks as described above.

**Lentiviral transduction of glioblastoma stem cells**

Scrambled and EMP3 shRNAs in lentiviral pTRIPZ expression vectors were transduced into NCH1425 and NCH644 GSCs by the DKFZ Cellular Tools Core Facility. Lentiviral particles were produced using a standard protocol. Briefly, HEK293FT (ThermoFisher, Braunschweig) cells were co-transfected with pTRIPZ-EMP3 shRNAs expression vectors and 2$^{nd}$ generation viral packaging plasmids VSV.G (Addgene #14888) and psPAX2 (Addgene #12260). Two days after transfection, viral supernatants were collected and filtered through a 0.45 µm filter. For lentiviral transduction $2.5 \times 10^6$ target cells (NCH1425 and NCH644 GSCs) were transduced by spinoculation (30 min; 800 g; RT) in the presence of 10µg/ml polybrene (Merck, Darmstadt). The resulting pellet was resuspended in complete growth medium and incubated for an additional 24 hours. After viral clearance, transduced cells were selected with 0.5 – 1.0 µg/mL puromycin (MP Biomedicals 194539). Knockdown efficiencies were validated by

Western blotting of lysates collected from cells after a 9-day induction period with 2 µg/mL doxycycline (MP Biomedicals 195044) administered at Days 0, 3, and 6 after seeding.

**Western blotting**

Total cell lysates were collected by scraping in NP-40 lysis buffer (50 mM Tris-HCl, pH 7.8, 150 mM NaCl, 1 mM EDTA, and 0.2% NP-40) supplemented with 1X Halt™ Protease and Phosphatase Inhibitor cocktail (Thermo Scientific 78440). Lysates were then snap-frozen in liquid nitrogen and thawed three times, and then centrifuged at 20,000 x g for 5 minutes. The clarified supernatant was collected and protein concentrations were measured by bicinchoninic assay (BCA). For each SDS-PAGE run, equivalent amounts of lysates in the range of 15-20 µg were prepared in 20 µL volumes containing 1X NuPAGE™ LDS Sample Buffer (Invitrogen NP0007) and 1X NuPAGE™ Sample Reducing Agent (Invitrogen NP0009). Lysates in reducing sample buffer were then heated at 95°C for 5 minutes, cooled to 4°C, and loaded into each well of NuPAGE™ 4-12% Bis-Tris gels (Invitrogen NP0322BOX) in NuPAGE™ MOPS SDS running buffer (Invitrogen NP0001). After the SDS-PAGE run (200 V constant for 1 hour), proteins were then transferred into a 0.45 µM nitrocellulose membrane (Invitrogen LC2001) using the XCell II™ Blot Module (Invitrogen EI9051) (30 V, 170 mA constant, 90 mins) and blocked with 5% milk in TBS-T for 1 hour. Membranes were then incubated with primary antibodies overnight at the appropriate dilutions (see Supplementary Information Table 2). The next day, membranes were washed thrice in TBS-T and incubated for 1 hour in the secondary antibody solution in 5% milk in TBS-T. Membranes were developed using WesternBright Sirius Chemiluminescent Detection Kit (Advansta K-12043-D20) and imaged using the chemiluminescence module of the Azure c400 Gel Imaging System (Azure Biosystems).

**Sample preparation for BioID2**

GBM cells stably transfected with the BioID2 bait constructs were seeded in triplicates at a density of 1.0 x $10^6$ cells in 10 mL of DMEM maintenance medium. Cells were then incubated at 37°C, 5% $CO_2$ for 48 hours until 70-80% confluence is reached. To induce biotinylation, old media was discarded and replaced with 10 mL of the same media supplemented with 50 µM biotin (Sigma-Aldrich B4639). Biotin-treated cells were further incubated at 37°C, 5% $CO_2$ for 18 hours. Cells were then lysed in 500 µL ice-cold NP-40 lysis buffer and lysates were further processed as described above.

To perform streptavidin pull-downs, 1 mL spin columns were washed with 200 µL of 1X PBS and packed with 100 µL of High Capacity Streptavidin Agarose Resin slurry (Thermo Fisher Scientific 20357). The columns were then centrifuged at 500 x g for 1 minute to remove the storage solution and then washed with 250 µL of 1X PBS thrice. A total of 1 mg of each lysate at 2 mg/mL concentration was then loaded into each column and incubated at room temperature for 30 minutes with end-over-end mixing. Columns were then centrifuged at 500 x g for 1 minute and washed four times with 250 µL of 1X PBS. Biotinylated proteins were then eluted by the addition of 2X reducing sample buffer (0.12 M Tris-HCl, 2% SDS, 20% glycerol, 2X lane marker tracking dye, pH 6.8) with 2 mM biotin and subsequent heating at 95ºC for 5 minutes. Eluates were collected by centrifugation at 1000 x g for 1 minute. Protein digestion for the BioID samples was then performed via a tryptic in-gel digestion protocol. For this, proteins were run for 0.5 cm into an SDS gel. After Coomassie staining, the total sample was cut out and digested with trypsin as previously described [6], adapted to on a DigestPro MSi robotic system (INTAVIS Bioanalytical Instruments AG, Cologne, Germany).

**BioID2 LC-MS/MS analysis**

Analysis was carried out on an Ultimate 3000 UPLC system (Thermo Fisher Scientific, Massachusetts, USA) connected to an Orbitrap Exploris 480 mass spectrometer (Thermo Fisher Scientific). Total LC-MS/MS analysis time was 90 min per sample. Prior to the analytical separation, peptides were online desalted on a trapping cartridge (Acclaim PepMap300 C18, 5 µm, 300 Å pore; Thermo Fisher Scientific) for 3 min using 30 µL/min flow of 0.05% TFA in water. The analytical multistep gradient was carried out on a nanoEase MZ Peptide analytical column (300 Å, 1.7 µm, 75 µm x 200 mm, Waters) using solvent A (0.1% formic acid in water) and solvent B (0.1% formic acid in acetonitrile). The concentration of solvent B was linearly ramped from 2% to 30% in 72 min, followed by a quick ramp up to 78 % B. After 2 min, the concentration of solvent B was lowered back to 2 % and a 10 min equilibration step appended. Eluting peptides were analyzed in the mass spectrometer using data-dependent acquisition (DDA) mode. A full scan at 60k resolution (380-1400 m/z, 300% AGC target, 45 ms maxIT) was followed by 1.5 sec of MS/MS scans. Peptide features were isolated with a window of 1.4 m/z and fragmented using 26% NCE. Fragment spectra were recorded at 15k resolution (100% AGC target, 54 ms. Unassigned and singly charged eluting features were excluded from fragmentation and dynamic exclusion was set to 35 s.

**Phosphoproteome and full proteome LC-MS/MS analysis**

LC-MS/MS analysis was carried out on an Ultimate 3000 UPLC system (Thermo Fisher Scientific) directly connected to an Orbitrap Exploris 480 mass spectrometer. Peptides were online desalted on a trapping cartridge (Acclaim PepMap300 C18, 5µm, 300 Å wide pore; Thermo Fisher Scientific) for 3 min using 30 µL/min flow of 0.05% TFA in water. The analytical multistep gradient was carried out using a nanoEase MZ Peptide analytical column (300Å, 1.7 µm, 75 µm x 200 mm, Waters) using solvent A (0.1% formic acid in water) and solvent B (0.1% formic acid in acetonitrile). A total of 150 min of LC-MS/MS analysis time was used per sample. The analytical step of the gradient was 134 min, during this time the concentration of B was linearly ramped from 4% to 30% (2%-28%, for phospho-peptides), followed by a quick ramp to 78%, after two minutes the concentration of B was lowered to 4% (2%) and a 10-min equilibration step appended. Eluting peptides were analyzed by the mass spectrometer using data-dependent acquisition (DDA) mode. A full scan at 120k resolution (380-1400 m/z, 300% AGC target, 45 ms maxIT) was followed by up to 1 s (2 s) of MS/MS scans. Peptide features were isolated with a window of 1.4 m/z (1.2 m/z), fragmented using 26% NCE (28% NCE). Fragment spectra were recorded at 15k resolution (100% AGC target, 22 ms maxIT; 200% AGC target, 54 ms maxIT). Unassigned and singly charged eluting features were excluded from fragmentation and dynamic exclusion was set to 35 s (10 s).

**Proteomics data analysis**

Data analysis was carried out via MaxQuant version 1.6.14.0 [7] using an organism-specific database extracted from Uniprot.org (For BioID2, a HUMAN reference proteome database containing 93267 unique entries from 01 August 2019, including the BioID2 sequences; for phosphoproteomics a HUMAN reference proteome database containing 79038 entries, from 03 January 2022) under default settings. Identification FDR cutoffs were 0.01 on peptide level and 0.01 on protein level. Match between runs (MBR) option was enabled to transfer peptide identifications across all RAW files based on accurate retention time and m/z for the BioID2. For the phosphoproteomics experiment MBR was restricted to only transfer peptide identifications within conditions. Raw files containing the full and phosphoproteomics data were analysed in separate parameter groups. For the phospho-fractions PTM was set to True and “Phospho (STY)” was added to the variable modification search. For the full proteome data and the BioID2 experiment, quantification was carried out using the label free quantification approach based on the MaxLFQ algorithm [2]. A minimum of 2 quantified peptides per protein was required for protein quantification.

**Statistical analysis and network visualization of BioID2 data**

BioID2 MaxQuant output files were further statistically analyzed using Perseus version 1.6.14.0 [8]. Briefly, potential contaminants and proteins that were not detected in at least one replicate of one experimental group were filtered out. Label-free quantification (LFQ) intensities were log2-transformed, and a two-sample, two-sided Welch's t-test was performed for each pairwise comparison between a bait (EMP3 WT or EMP3 N47A) and a spatial reference control (TagRFP or GAP-TagRFP). Proteins with a difference in log2(LFQ) ≥ 1 and p-value ≤ 0.05 were considered significantly enriched in bait pull-downs. To identify proteins that were uniquely interacting with the EMP3 WT or EMP3 N47A bait protein, additional filtering was performed for hits that were present in all EMP3 WT/N47A replicates but absent in all replicates of either spatial reference control pull-downs. To map the proteins into a network, protein lists were submitted to STRING (version 11.5) and the resulting network exported into Cytoscape (version 3.9.1). Additional filtering for edges with STRING scores > 0.700 (i.e., high confidence interactions) and nodes with degrees ≥ 3 was performed. Enrichment analysis was performed using the enrichment analysis plug-in available in the stringAPP (version 1.7.1). Nodes were colored and organized according to the functional groups defined by GO or Kyoto Encyclopedia of Genes and Genomes (KEGG) terms, while plasma membrane localization scores based on the COMPARTMENTS database [1] were continuously color-mapped onto the node edges. Dot plots depicting enrichment scores and significance values of proteins from each major cluster were further generated using ProHits-viz [3].

**Kinase enrichment analyses**

MaxQuant output files were statistically analyzed using Perseus version 1.6.14.0 [8]. First, LFQ intensities were log2-transformed. Potential contaminants and phosphosites containing invalid values (i.e., LFQ = 0) in more than 2 replicates per experimental group were filtered out and not considered in the downstream analysis. In addition, only phosphosites with localization probabilities ≥ 0.75 were retained. The remaining missing values were imputed separately for each replicate using values from a normal distribution (width = 0.3, down shift = 1.8). Afterwards, a two-sample, two-sided Welch's t-test was performed comparing control and EMP3 KO conditions within each cell line. Phosphosites with

|log2-FC|≥ 1 and FDR-adjusted p-values ≤ were considered as differentially phosphorylated when comparing EMP3 KOs to their respective controls.

To perform phosphorylation analysis in IPA, the input list of differentially phosphorylated proteins along with their respective phosphorylation log2-FCs and affected phoshosites were imported into IPA version 22.0 (Qiagen). Default parameters for phosphorylation analysis were used, except for the Species parameter which was restricted to "Human" only. For kinase enrichment analysis (KEA), proteins with commonly dephosphorylated sites (i.e., dephosphorylated in both DK-MG and U-118 EMP3 KOs compared to their corresponding controls) were imported into KEA version 3 (KEA3; https://maayanlab.cloud/kea3/, accessed 04 August 2022). The top 5 upstream kinases based on FDR-adjusted p-values were identified using the KSIN, PhosD, and PTMsigDB kinase-substrate databases as reference. For Robust Inference of Kinase Activity, phosphosites that were commonly regulated in DK-MG and U-118 EMP3 KOs (i.e., phosphosites with log2-FCs in the same direction) were imported into RoKAI version 2.1.3 (https://rokai.io/, accessed 04 August 2022) along with their respective log2-FC values. Top 10 upstream kinases based on FDR-adjusted p-values were then identified. For all analyses, the R package ggplot2 was used to visualize identified upstream regulators along with their respective significance levels and associated substrates.

**Quantitative reverse transcription PCR**

Total RNA were extracted in three biological replicates from DK-MG and U-118 control and EMP3 KO cells using the Nucleospin RNA kit (Macherey-Nagel) following the manufacturer's instructions. For each extraction, the equivalent cDNA was then synthesized from 1 µg of RNA using the Superscript IV First Strand Synthesis System (Thermo Fisher Scientific 18091050) with 2.5 µM oligo-dTs and 2.5 µM random hexamers as primers. Possible genomic DNA contamination was assessed by adding parallel minus RT reactions. qPCR was then performed on a CFX96 qPCR Real-Time PCR module with C1000 Touch Thermal Cycler (Bio-Rad) using Quantitect SYBR Green PCR Master Mix (Qiagen) and Quantitect Primer Assays (Qiagen) for CDC6, MCM7, ORC6, RFC2, and GAPDH. The following program was used for the qPCR run: 1) initial activation at 95°C for 5 minutes, followed by 40 cycles of 2) denaturation at 94°C for 15 seconds, 3) annealing at 55°C for 30 seconds, and 4) extension at 72°C for 30 seconds. Relative expression levels were then quantified based on Ct values using GAPDH as the reference target. Unpaired one-tailed t-tests were then performed to confirm that the selected DEGs are significantly reduced in EMP3 KOs versus their respective controls.

**Immunofluorescence**

Cells were seeded at a density of 1.0 x $10^4$ cells/mL of DMEM maintenance media in 4-well chamber slides and incubated at 37°C, 5% $CO_2$ for 72 hours. To perform immunostaining, cells were washed once with 1X PBS and then fixed with 4% formaldehyde (Carl-Roth 4980.2) in 1X PBS at room temperature for 15 minutes. Each well was then washed with 1X PBS for 5 minutes thrice, and fixed cells were blocked with 0.1% saponin, 5% FBS in 1X PBS under gentle shaking at room temperature for 1 hour. Cells were incubated with the primary antibodies at the appropriate dilutions (Supplementary Table 2) in blocking buffer at 4°C overnight. The next day, the primary antibody solution was discarded and cells were washed again thrice in 1X PBS. Afterwards, primary antibody-stained cells were incubated at room temperature in the dark with the appropriate secondary antibodies diluted in blocking buffer. After three washes with 1X PBS, slides were dried and mounted using Vectashield® HardSet™ with DAPI (Vector Laboratories VEC-H-1500). Images were then captured with Leica TCS SP5 II confocal microscope (Leica Microsystems, Wetzlar, Germany) using 63X/1.4 NA objective and the appropriate laser lines and filters.

**CellTiter-Glo assay**

To measure daily proliferation rates of control and EMP3 KO U-118 and DK-MG cells, 2.0 x $10^4$ cells were seeded in 100 µL of DMEM maintenance medium in each well of a white, flat-bottom 96-well plate. A total of 5 identical plates were prepared, each corresponding to the daily time point starting from Day 0 to Day 4 post-seeding. To measure the mitogenic response of the same cell lines to EGF, 10,000 cells were seeded in 500 µL of DMEM maintenance medium in a 24-well plate and incubated overnight at 37°C, 5% $CO_2$. The next day, cells were serum-starved by replacing DMEM maintenance medium (supplemented with 10% FBS) with DMEM only. Serum-starved cells were then treated with 100 ng/mL EGF daily for 3 days, and the number of viable (i.e., metabolically active) cells of treated and untreated control and EMP3 KO cells were measured one day after the last EGF treatment. To measure the effect of AZD9291 on cell viability, cells were also seeded in a white 96-well plate as described in the proliferation assay. After a 24-hour incubation at 37°C, 5% $CO_2$, cells were treated with increasing concentrations of AZD9291 (Hölzel Diagnostika, HY-15772-10 mM) and further incubated for another 24 hours. For all three experiments, 100 µL of CellTiter-Glo 3D (Promega G9683) were then added to each well at the time of measurement. After shaking at 300 rpm for 2 minutes, luminescence levels were measured using the FLUOstar Omega microplate reader (BMG Labtech, Ortenberg, Germany) coupled to the Omega software (version 5.11 R4). Relative luminescence units (RLU) were calculated by normalizing to the appropriate controls and plotted and statistically analyzed in GraphPad Prism (version 9.3.1).

**Caspase 3/7 and PARP cleavage assay**

GBM cells were seeded at a density of 10,000/200,000 cells in 100 µL/2 mL of DMEM maintenance medium per well for the Caspase 3/7 and PARP cleavage assays, respectively. After a 24-hour incubation at 37°C, 5% $CO_2$, apoptosis was induced by treating the cells with 100 nM and/or 1 µM staurosporine (STS; Sigma-Aldrich 569396) for 4 hours or 2.5 µM AZD9291 for an additional 24 hours. Caspase 3/7 levels were measured using the Caspase-Glo 3/7 Assay Kit (Promega G8091). Briefly, 100 µL of Caspase 3/7 reagent were added to each well. The plate was mixed at 500 rpm for 30 seconds. After a 30-min incubation time at room temperature, luminescence levels were recorded using the microplate reader. RLU were calculated by normalizing to untreated cells, and the results were plotted and statistically analyzed using GraphPad Prism (version 9.3.1). To assess extent of STS-induced PARP cleavage, total cell lysates were collected, and Western blotting was performed as described above.

**Treatment of GSCs with CDK2 inhibitor**

NCH1425 and NCH644 GSCs were seeded at a density of 750,000 cells in 6 mL of GSC maintenance medium containing DMSO (Sigma-Aldrich D2650) or doxycycline (MP Biomedicals 195044) at a concentration of 2 µg/mL (Day 0). At Day 3 post-seeding, induced cells were treated again with DMSO or doxycycline and incubated at 37°C, 5% CO2 for another 3 days. At Day 6, cells were split, re-induced or treated with DMSO, and re-seeded into 96-well white flat bottom plates (Corning 353296) at a density of 5,000 cells per well in 90 µL of GSC media. After 24 hours of incubation, cells were treated with increased concentrations of the CDK2-specific inhibitor K03861 (Hölzel Diagnostika A15889) dissolved in 10 µL of GSC media. After a 24-hour incubation, cell viabilities were measured using the CellTiter-Glo assay as described above. For each condition, percentage changes in cell viabilities as a result of drug treatment were calculated by normalizing the CTG values of K03861-treated cells to untreated controls (set as 100%). The fold-change in viability due to induction of shRNA expression at each concentration was then calculated by normalizing percentage changes of induced cells to uninduced cells. Results were then plotted and statistically analyzed in GraphPad Prism (version 9.3.1).
